# Time-calibrated genomic evolution of a monomorphic bacterium during its establishment as an endemic crop pathogen

**DOI:** 10.1101/2020.05.26.115717

**Authors:** Damien Richard, Olivier Pruvost, François Balloux, Claudine Boyer, Adrien Rieux, Pierre Lefeuvre

**Author notes:** Corresponding author: Damien Richard. **Competing interests:** The authors declare that they have no competing interests.

## Abstract

The reconstruction of the evolutionary histories of pathogen populations in space and time has greatly improved our understanding of their epidemiology. However, analyses are usually restricted to the non-recombining genomic regions and, thus, fail to inform on the dynamics of the accessory genome. Yet, horizontal gene transfer is of striking importance to the evolution of bacteria as it can redistribute phenotypically important genes. For bacterial pathogens, those include resistance to antimicrobial compounds and virulence factors. Understanding the gene turnover in genomes at microevolutionary scales is key to apprehend the pace of this evolutionary process. Here we addressed this question for the epidemic lineage of a major bacterial plant pathogen, relying on a dense geographic sampling spanning 39 years of evolution. Gene turnover rate exceeded SNP mutation rates by three orders of magnitude. Accessory genes were preferentially plasmid-encoded, but we evidenced a highly plastic chromosomal region hosting ecologically important genes such as transcription activator-like effectors. We argue that turnover of accessory genes provides a potent evolutionary force in monomorphic bacteria, and exemplify this statement retracing the history of a mobile element conferring resistance to copper compounds widely used for the management of plant bacterial pathogens.

## Introduction

Recently, the retrospective analysis of time and geographically informed genomic datasets has gained in popularity for reconstructing the evolutionary history of pathogens (1). It can shed light on the pathogens’ origin and transmission pathways, as well as on the evolution of key phenotypes (2, 3). In addition to time calibrating the pathogens’ evolution from bacterial core-genome kinship, the analysis of genomic datasets can reveal the importance of gene content variation relative to nucleotide substitution. As such, multiple genome-wide studies evidenced large gene content variation and the importance of recombination as a major evolutionary force in many bacterial species (4–8). Gain and loss of accessory genes is primarily achieved through plasmid or bacteriophage acquisition (9–11) and plays an important role in bacterial genomic evolution (5). Indeed, accessory genes can provide access to novel functions, such as enhanced in-host multiplication, the ability to colonize a new ecological niche or antimicrobial resistance and thus can improve the fitness of a bacterial lineage (12–14). Surprisingly, studies to determine the rate of gene content variations are still scarce (15–19). The concomitant reconstruction of ancestral gene content in a combined approach has recently shown promise for deciphering transmission, geographical expansion and host specificity for the study of plant pathogenic bacteria (20–22). This approach appears particularly relevant for the study of bacterial lineages that display low SNP substitution rates (referred to as monomorphic (10)).

*Xanthomonas citri* pv. *citri* (*Xcc*), a gamma-proteobacterium that originated from Asia and subsequently spread to other regions, is the causal agent of the Asiatic citrus canker, an economically important disease that threatens citrus industries in most areas of production. *Xcc* is a monomorphic bacterium, whose genome was reported to display variations through differences in genomic island composition (23) or plasmid content (24, 25). Indeed, some key phenotypes are encoded by plasmid genes: streptomycin resistance (26), copper resistance (24, 27) and most importantly type III transcription-activator like effector (TALE) genes (28). *Xcc* TALEs include a large repertoire of related but highly variable genes, of which *pthA4* is required for pathogenicity (29).

Herein, using a comprehensive collection of 221 genetically-related *Xcc* strains collected in the South West Indian Ocean (SWIO) region over a 39-year timespan, we compared the gene and nucleotide substitution rates in order to quantify their respective effect on genome evolution. Besides time calibrating and quantifying *Xcc* evolution, we put forward hypotheses on the long-term history of *Xcc* in the SWIO region, going back to its first probable introduction ca. two centuries ago. We highlighted a very high ratio of gene content changes to mutation, emphasized that most gene content changes are short-lived, and evidenced that most accessory genes located on plasmids and on a newly described chromosomal genomic island.

## Results

### Global phylogeny

The analysis of 284 strains representing worldwide *Xcc* diversity with a particular emphasis on the SWIO islands (Figure 1) revealed 7 005 high confidence SNPs. The phylogenetic reconstruction obtained from this set displayed a strong geographic structure (Supplementary Figure S1). Whilst some subclades comprising worldwide strains remained unresolved, all 210 strains sampled from the SWIO region formed a well-resolved monophyletic clade within which subclades tend to group strains sampled from a single island (Figure 2). No obvious spatial structure was apparent within Réunion and all but one of the citrus groves hosted strains from multiple Réunion subclades (Supplementary Figure S1B). Consistent with previous data (20), copper-resistant (Cu^R^) *Xcc* from Martinique (French West Indies) were closely related to most Cu^R^ strains isolated from Réunion.

**Figure 1.**
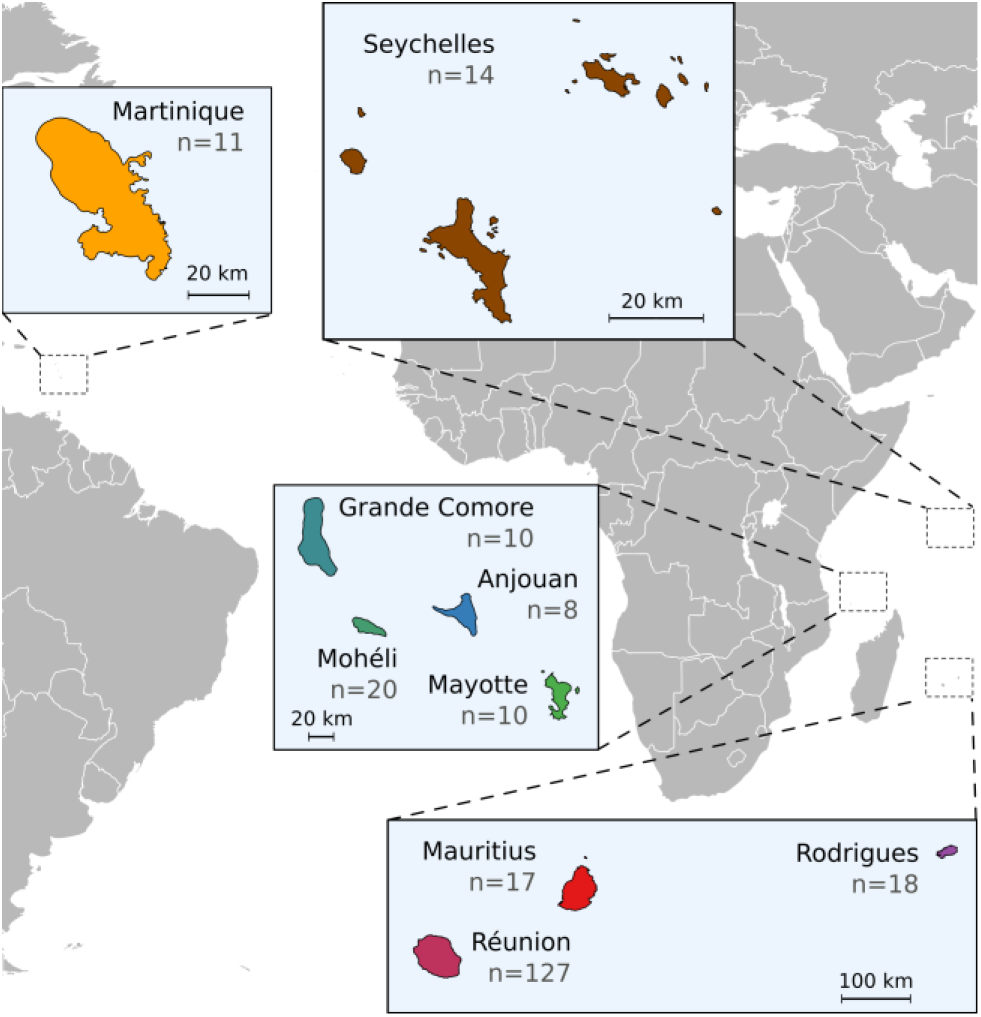
Location of the South West Indian Ocean islands where the majority of the strains studied were sampled. Number of strains per location is indicated in parentheses.

**Figure 2.**
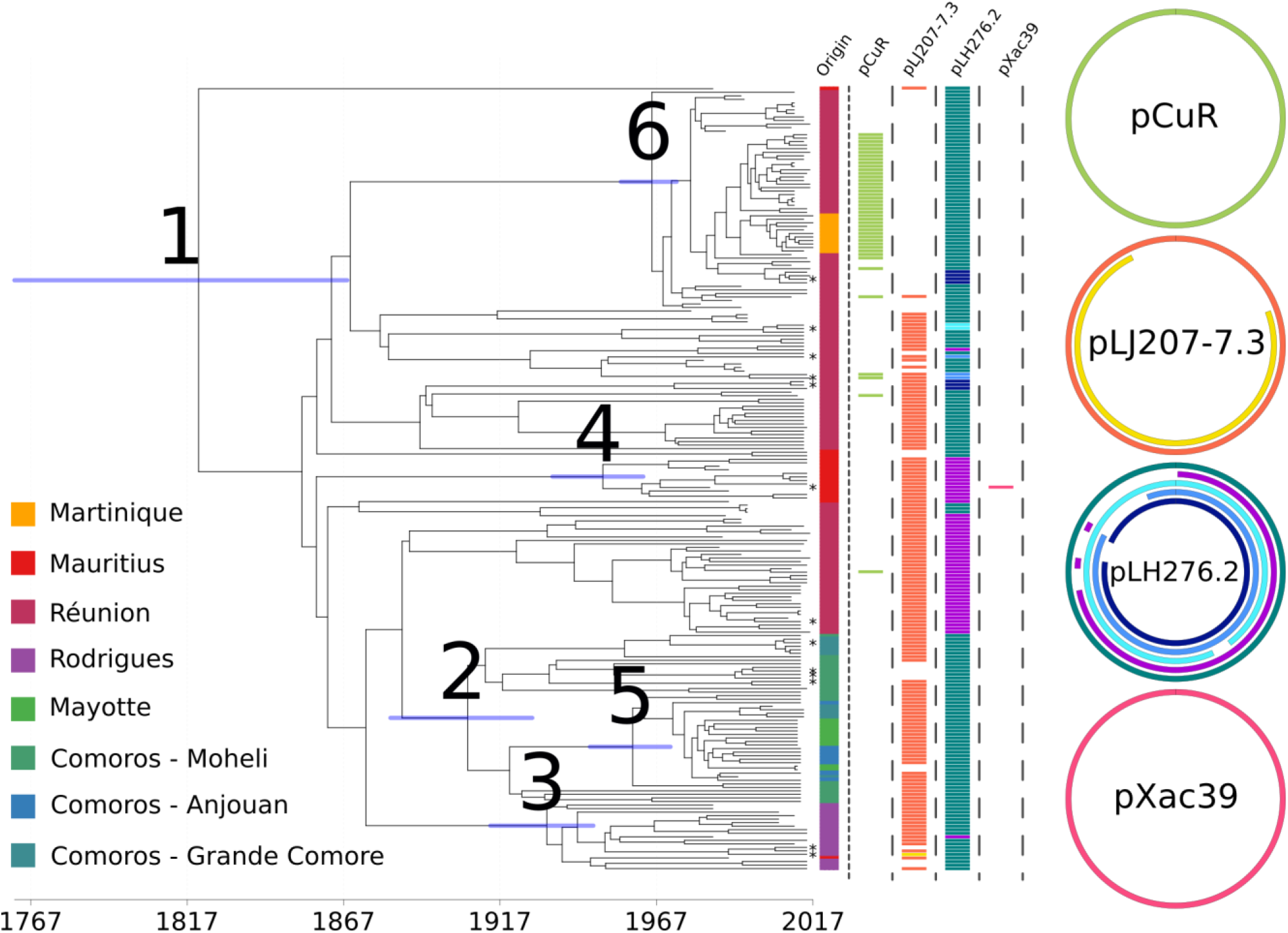
A dated phylogeny of the SWIO clade of *Xanthomonas citri* pv. *citri*. X-axis under the phylogenetic tree represents the timescale in years (AD). Node bars represent 95% highest posterior density (HPD) for node ages estimated with tip-calibration. Tips are coloured according to sampling location. Coloured boxes indicate the plasmid content of each strain. The colour code for the boxes matches the colours for the represented circular plasmids and corresponds to observed plasmid alleles. Node numbers correspond to those in Table 1. Tip labels with asterisks indicate strains for which long-read sequencing was also performed.

### Temporal signal and time calibration

Although there was no significant temporal signal for the global phylogeny, the regression between the root-to-tip distance and node age was significant (*p*-value 2.3×10^−4^ and R^2^ 0.06) at the root of the SWIO clade with a slope of 3.2×10^−5^ (Supplementary Figure S2B). The temporal signal for the SWIO clade was further confirmed by a date-randomization test (Supplementary Figure S2A).

**Table 1.** Inferred dates of MRCA and substitution rates of the SWIO clade along with those of five clades of interest.

| Node number | Inferred node date | Date 95% confidence interval | No. of strains | No. of variable genes | No. of variable SNPs | Subst. rate SNP | No. of core-genome genes | No. of pan-genome genes |
| --- | --- | --- | --- | --- | --- | --- | --- | --- |
| 1 | 1818 | 1762-1868 | 221 | 699 | 3403 | 8.8E-8 | 4347 | 5046 |
| 2 | 1906 | 1882-1927 | 48 | 205 | 812 | 8.8E-8 | 4546 | 4751 |
| 3 | 1931 | 1914-1947 | 19 | 222 | 363 | 8.8E-8 | 4531 | 4753 |
| 4 | 1949 | 1934-1963 | 12 | 143 | 134 | 8.5E-8 | 4611 | 4754 |
| 5 | 1959 | 1946-1971 | 27 | 147 | 288 | 8.3E-8 | 4593 | 4740 |
| 6 | 1965 | 1956-1973 | 62 | 358 | 318 | 8.1E-8 | 4599 | 4957 |

A time-calibrated analysis was conducted using the BEAST framework (30) on the 221 strains of the SWIO clade, using an exponential growth tree model and an uncorrelated lognormal relaxed clock (estimated standard deviation: 0.26 (95% HPD 0.18-0.34)). Overall, the Maximum Likelihood and the Bayesian trees had a similar topology (Supplementary Figure S3). The SWIO tree root age was established at around 1818 (95% HPD: 1762-1868, Table 1), with an estimated substitution rate of 8.4×10^−8^ substitutions per site per year (95% HPD: 6.9×10^−8^-1.0×10^−7^). This corresponds to 0.43 SNP substitutions per genome per year (95% HPD: 0.35-0.51).

### Accessory genome content

In order to test our ability to reconstruct the gene content of *Xcc* strains, two strains were sequenced in triplicate before the application of the gene content inference pipeline. Gene content (4903 genes) was identical between all replicates for one of the strains, while six genes upon 4847 were variably detected from the second strain replicates (replicates pairwise maximum difference of four genes). Importantly the variably detected genes were not clustered within the genomes. It is therefore likely that events (i) involving the gain or loss of numerous genes or (ii) displaying a phylogenetic signal (i.e. when the same event is revealed in closely related individuals) are genuine.

After *de novo* genome assembly, gene detection and gene clustering of the 221 SWIO strains, a pan-genome size of 5 046 genes was identified. The core genome comprised 4 347 genes (i.e. detected in all strains), while the accessory genome comprised 699 genes (i.e. absent from at least one strain). Of these accessory genes, 336 were assigned to the chromosome, 339 to plasmids and 24 remained unassigned (Figure 3). As shown by the gene content analysis of sequencing replicates, our gene detection pipeline may have limitations associated with the detection of unique genes that have been gained or lost in single strains. Fourteen out of 699 genes displayed such characteristics. Most of the accessory genes (70%) did not match any known Cluster of Orthologous Group (COG); 4% matched replication, recombination and repair functions; 3% inorganic ion transport and other COG categories each comprised fewer than 3% of the gene clusters. We then used a Bayesian reconstruction of the ancestral states in terms of gene presence/absence to estimate gene gain, loss and total substitution rates for both the plasmid and the chromosome compartments. The plasmid gene substitution rate (1.75×10^−3^ gene substitutions per gene and per year 95% HPD: 1.40×10^−3^-2.96×10^−3^) appeared to be significantly higher than the chromosome gene substitution rate (7.00×10^−5^, 95% HPD: 6.57×10^−5^–8.21×10^−5^). Nevertheless, at the replicon scale, these rates largely overlapped with 0.27 gene substitutions per genome and per year for the plasmid compartment (95% HPD 0.22–0.46) and 0.32 for the chromosomal compartment (95% HPD 0.30–0.38). Chromosome gene gain and loss rates were in the same order of magnitude and showed overlapping 95% HPD (gene gain rate 1.06×10^−5^, 95% HPD 7.75×10^−6^-1.55×10^−5^ and gene loss rate 5.94×10^−5^, 95% HPD 5.70×10^−5^-6.61×10^−5^, respectively). Similar results were observed for the plasmid compartment (gene gain rate 1.12×10^−3^, 95% HPD 9.21×10^−4^-1.68×10^−3^ and gene loss rate 6.27×10^−4^, 95% HPD 3.98×10^−4^-1.22×10^−3^).

**Figure 3.**
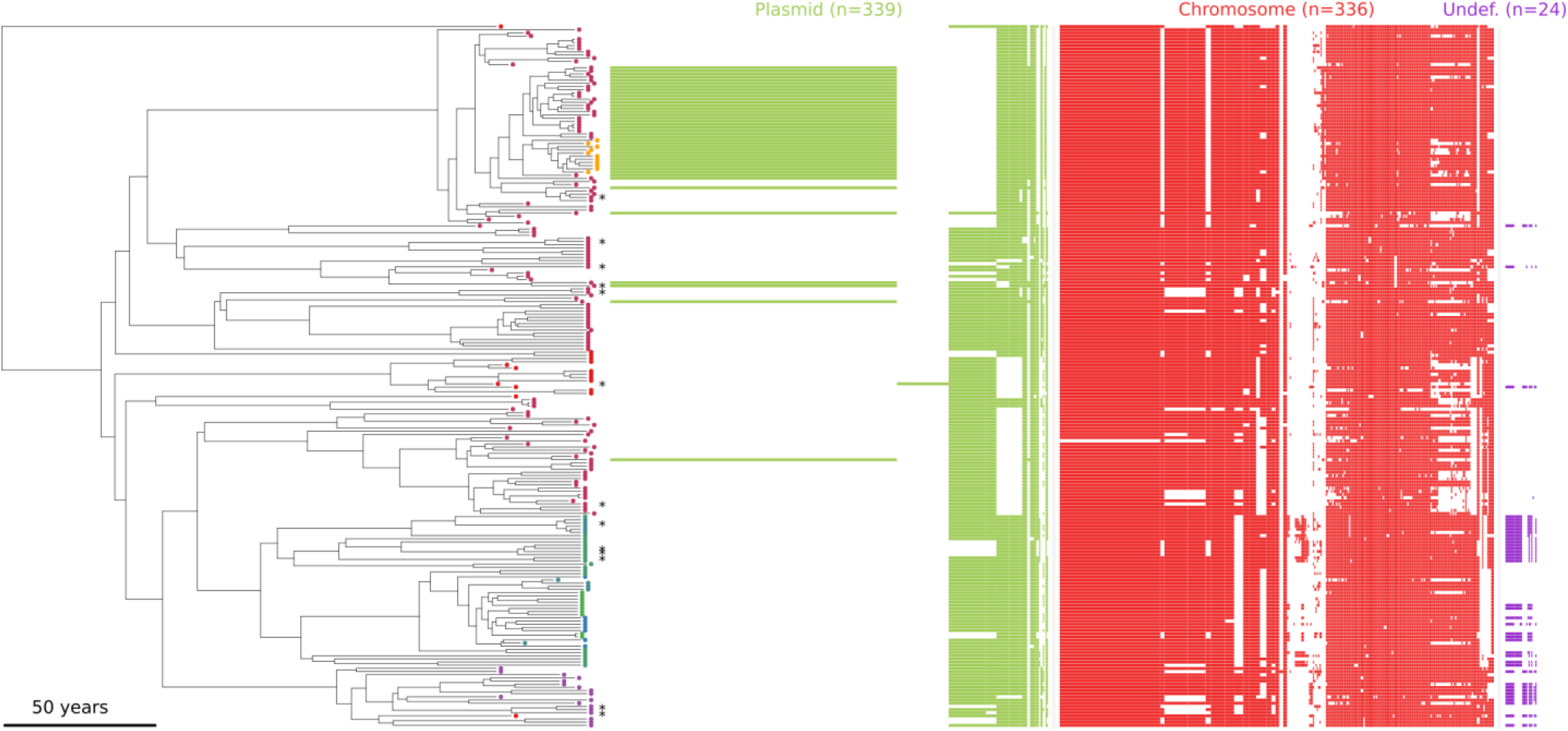
Time-calibrated phylogeny of the SWIO clade. The matrix on the right of the tree comprises one column for each of the 699 genes that varied in presence/absence among the 221 SWIO strains. A coloured box is printed in front of a strain if it encodes for this gene cluster. Gene clusters are separated according to their genetic compartment of origin: plasmid (green), chromosome (red) or undefined (purple). Tips with asterisks indicate strains for which long-read sequencing was also performed.

### Gain, loss and mosaic structure of plasmids

Whereas it was not possible to achieve full plasmid reconstruction using our short-read and gene-based approach, we were able to assess the presence of the gene content associated to the known set of *Xcc* plasmids (31) in each strain. We were able to define the presence of four distinct plasmid groups with up to five distinct alleles associated with the SWIO *Xcc* strains (see the right side of Figure 2). Importantly, these alleles were confirmed with long-read sequencing and assembly on a subset of strains (strains indicated with an asterisk on Figure 2, Supplementary Table S1).

Asiatic citrus canker control largely relies on the repeated applications of copper-based pesticides (32). In Réunion over the last decade, copper resistance has emerged with the integration from an unknown source of a mobile heavy metal resistance plasmid (pCu^R^) in previously established *Xcc* strains (24). Herein, we observed that the copper-resistant phenotype was always associated to the presence of pCu^R^. Interestingly, the pCu^R^ gene content was identical for its 42 occurrences (31 in Réunion and 11 in Martinique). We inferred that pCu^R^ was gained 6.9 times and lost 0.84 times (95% HPDs 6-9 and 0-3, respectively) in the phylogeny. Thirty-four filtrated SNPs were detected within pCu^R^ sequences, but with no significant temporal signal (data not shown). Most of the pCu^R^-bearing strains (n=37) formed a monophyletic group within subclade 6, the remaining strains (n=5), originating from Réunion, were distributed in three distinct subclades (Figure 2). Fully identical pCu^R^ plasmids (without any SNP) were carried by distantly related strains isolated both in citrus groves and a commercial nursery (Supplementary Figure S4).

In contrast with the dispensable character of pCu^R^, all *Xcc* strains known to date encode a pathogenicity-related plasmid gene set, organized as one or multiple plasmids (a few of which have been described so far (25, 28).

*Xcc* pXac64-like plasmids (28) typically carry *pthA4*, a transcription activator-like effector (TALE) gene required to produce citrus canker symptoms (14). As expected, a pXac64-like plasmid was found in every SWIO strain. Five distinct plasmid variants were detected. In particular, 49 strains from Mauritius, Réunion and Rodrigues displayed a deletion of 26 genes mostly encoding for plasmid transfer and maintenance functions (Figure 2).

The plasmid pLJ207-7.3 is closely related to pXac47 (33) but comprises a TALE gene whereas pXac47 doesn’t. The functions of the proteins encoded by these plasmids were mostly unknown (n=24), but included plasmid maintenance and transfer (n=17), as well as other functions (n=15). While most strains from all SWIO islands carried a pLJ207-7.3-like plasmid, it was absent in 77 SWIO strains (Figure 2). Of these, 61 were phylogenetically related and corresponded to all but one individual of clade 6, which included most of the Cu^R^ strains. Sixteen other strains lacking the plasmid were scattered throughout the phylogeny and had been sampled in Réunion, Mauritius, Mohéli, Mayotte and Rodrigues.

As for pCu^R^, both pXac64 and pXac47 variants were scattered along the phylogeny. The geographic distribution of the plasmid alleles did not display any clear structure. Indeed, we observed up to six distinct plasmid profiles in a single grove in Réunion.

Finally, a previously unreported 39.8kb plasmid encoding 40 genes was found in a single strain from Mauritius. The existence of the plasmid was confirmed through long-read sequencing. No other strain presented these genes. Besides conjugation and plasmid partitioning, no specific function could be associated to the plasmid. Similar plasmids (nucleotide identity >75%) were previously found in *Xylella fastidiosa* or *Xanthomonas oryzae* (GenBank accession CP014330.2 and CP007810.1, respectively).

### A highly polymorphic chromosomal region

Compared to plasmids, *Xcc* chromosomal gene content was rather homogeneous: we only detected 336 chromosomal accessory genes. Using gene positions of a high-quality circular chromosome sequence (GenBank accession CP018854.1, from SWIO strain *Xcc* LH276) as a reference, we located 118 SWIO chromosomal accessory genes in one genomic island (GI). The remaining 218 genes were spread along the chromosome in groups of up to seven genes or had no homologue in the chosen reference. The GI did not match recombinant loci, as shown by our analysis and that of Gordon et al. (23). Long-read based *de novo* assemblies of 13 strains demonstrated that the chromosome of *Xcc* is apparently not rearrangement-prone (only one strain displayed a 1.2Mb inversion when compared to LH276 reference chromosome). It also confirmed the extensive variation in the GI’s gene content, which carried between 32 and 92 genes, bordered by a phage integrase and a complete IS*3* element on the 5’ side and by one or multiple incomplete *AttR* sites on the 3’ side (Figure 4). No deviation in GC content was detected for this region (data not shown). In most strains, the GI carried putative multidrug efflux pumps and a class C beta-lactamase was detected in five out of 13 strains. Most notably, the GI comprised one or two usually plasmid-borne TALE genes in seven strains (Figure 4). Chromosomal TALEs harboured between 8.5 and 28.5 repeats. Due to the repetitive nature of TALE genes, only long reads that covered the entire gene were used to generate the GI assemblies, therefore reducing the coverage of this region and impairing precise RVD motif determination.

**Figure 4.**
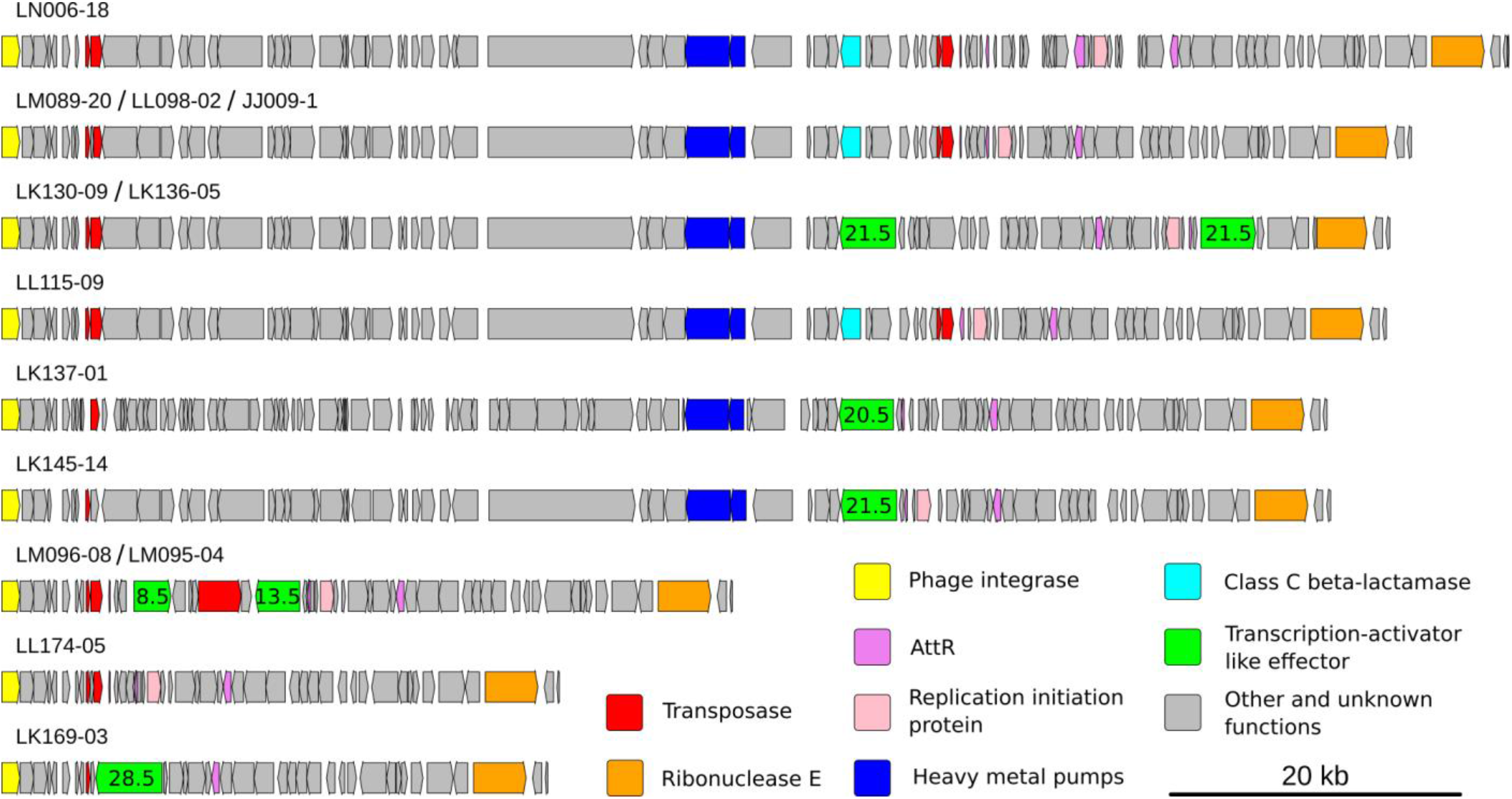
Representation of the gene content of the chromosomal genomic island (corresponding, in LH276, to positions 2 832 588 – 3 003 260) for 13 strains sequenced using long reads. Blocks represent genes homologous to known *Xanthomonas citri* pv. *citri* proteins and are coloured according to the predicted function of their encoded proteins (see legend). Number of Repeat Variable Di-residue of Transcription-Activator-Like Effector (TALE) genes, as estimated based on error-prone long reads, are written in the corresponding boxes.

## Discussion

In this study, we applied a population genomic approach on a set of bacterial strains obtained from an epidemic clade of *Xcc*, the causative agent of Asiatic citrus canker. We dated the introduction of *Xcc* in the region to 1762-1868 and were able to reconstruct its local evolutionary history in the SWIO islands. We conjointly inferred the SNP and gene substitution rates and evidenced that gain and loss of accessory genes in *Xcc* is pervasive, even at the most narrow spatiotemporal scale.

### Emergence of *Xcc* in the SWIO area

We used the age of the ancestor of all the strains as a proxy to determine the date when *Xcc* was introduced in the SWIO area. It was estimated to be 1818 (95% HPD: 1762-1868), which predates the earliest report of the disease in the area (1917 in Mauritius, (34)). *Xcc* and its main host genus, *Citrus*, originated in Asia (35, 36) and were probably spread beyond their area of origin by human-mediated movements of plants or plant propagative material. The timeframe suggested here yielded two main hypotheses regarding the origin of the pathogen in the SWIO.

The first hypothesis imply the introduction of contaminated plants by the French botanist and colonial administrator Pierre Poivre (1719-1786) thought to be at the root of the first introduction of citrus in the Mascarene Archipelago from several Asian countries in the mid-18^th^ century (37). The second hypothesis is that after the abolition of slavery in Mauritius and Réunion in 1835 and 1848, respectively, hundreds of thousands of indentured labourers from several Asian countries (the majority from India) were brought in to boost the agricultural workforce (38–40). This active flow of goods and people from the Asian continent may have led to the concomitant introduction of *Xcc* in the SWIO area. Indeed, the emergence of pre-adapted plant pathogens due to migratory events is a well-recognized phenomenon (41–43). We were unable to accurately identify the geographic origin of the strains that first migrated to the SWIO region. More strains from the hypothetical Asian cradle of *Xcc* will have to be sequenced to provide a definite answer regarding the pathogen’s geographic origin and migratory history.

### Genetic structure within SWIO

Contrasting with the genetic diversity found in the northern Indian Ocean and India, SWIO strains grouped in a single clade. This monophyly indicates that the strains first introduced in the area were closely related, genetically and probably geographically. It also suggests the absence of recent introductions from remote countries. We observed a strong geographic structure within the SWIO. However, Réunion and Mauritius strains were both polyphyletic, impairing the identification of the location where the pathogen first established. This polyphyly could originate from a polyclonal primary inoculum, a frequent case for *Xcc* (44). It could also evidence ancient exchanges of strains between islands. Indeed, the strain phylogeny also suggests recent inter-island migration events between (i) Mauritius and Rodrigues (a remote Mauritian territory), (ii) the four islands in the Comoros Archipelago and (iii) Réunion and Martinique. In all three cases, political and/or economic links between islands are in accordance with the usual long-range dispersal of *Xcc* through human-associated dispersal of contaminated citrus material (44).

In contrast with the strong geographic structure of *Xcc* between islands, the analysis of strain diversity obtained after dense sampling in Réunion revealed little concordance between phylogeny and grove location, possibly reflecting the multiplicity of inoculum inputs during the lifespan of citrus groves. While grove contamination can occur via infected nursery plants when the grove is first established, it also occurs during periods of wind and rain (*i.e.* storms or hurricanes regularly hit islands in the region) and grove maintenance operations (32).

In Réunion, severe *Xcc* outbreaks happened during weather conditions which were usually unfavourable to disease development. These were found to be related to the emergence of copper-resistant strains (45). The presence of fully identical pCu^R^ plasmids in phylogenetically distant strains further confirmed previous data and *in vitro* tests that evidenced pCu^R^’s mobility. Additionally, the loss rate of pCu^R^ was found to be seven times lower than its gain rate, suggesting that pCu^R^ is crucial to *Xcc* in the context of Réunion citriculture with repeated copper applications. Its stability can also reflect the functioning of plasmid maintenance systems such as toxin-antitoxin or partitioning. The gene content uniformity of pCu^R^ contrasts with its mosaic nature on a broader geographic and phylogenetic scale (24). This suggests that it was recently gained by *Xcc* (46) and has spread rapidly through commercial citrus nurseries.

### Genetic support of TALE genes

While *Xcc* chromosomal synteny and gene content were relatively homogeneous, a GI region located downstream to a phage integrase and carrying one or multiple partial copies of an *AttR* site in its 3’ end was highly variable in terms of gene content and organization. *AttR* and *AttL* motifs result from phage integration into a genome. The lack of the *AttL* motifs within the variable chromosomal region suggests the presence of a non-functional and/or remnant phage. Interestingly, some allelic versions of the GI comprised putatively adaptive genes, including a heavy metal efflux pump, an antibiotic resistance gene and up to two type III effectors of the TALE family. Bacterial TALE genes are crucial for pathogenicity. This family of genes codes for proteins that trigger the activation of specific plant genes and aid plant infection. While one of these genes (*pthA4*) is recognized as the major pathogenicity determinant of *Xcc* pathotype A, *Xcc* genomes always contain several alternative TALE genes, some of which were found to modify its virulence (47, 48). Until recently, TALE genes were thought to be only plasmid encoded in *Xcc*. A recent study reported a chromosomal TALE in *Xcc* strains with a restricted host range (known as *Xcc* pathotype A^w^) (49). In our study, we report a similar result for *Xcc* strains with a broad host range, including all citrus cultivars (pathotype A). The presence of insertion sequences of the IS*3* family, both in TALE-bearing plasmids and in the GI, may have mediated homologous recombination between plasmid and chromosome, as reported earlier in *Escherichia coli* (50). Additionally, *Tn3* family transposons might be involved both in plasmid rearrangement and TALE gene evolution in *Xcc* (25, 51). On *Xcc* plasmids, TnpA, the transposase required for the autonomy of *Tn3* transposons (52), is sometimes bordered by two TALE genes, forming a tripartite element that we found in the chromosomal GI of two strains (Figure 4).

### Rates of evolution: SNP and gene substitution rates

We estimated that *Xcc* SNP substitution rate comes within the lower range of known bacterial rates, in line with its monomorphic nature (53). Interestingly, when expressed on a per-site basis the gene loss and gain rate for was three orders of magnitude higher than SNP substitution rates. Some previous studies involving similar analyses reported that gene turnover rates were in the same order of magnitude as the SNP rate (15–18). It is important to note that these studies were performed at deeper phylogenetic scales than ours. Indeed, the gene substitution rate is highly dependent on the evolutionary timescale considered, given that most gene content variations are usually deleterious and short lived (7, 19, 54). Vos et al. (19) (i) highlighted a marked lack of available data on gene substitution rates in the literature, despite its upmost importance in bacterial genome evolution, and (ii) emphasized the importance of analysing datasets composed of closely-related strains in order to explicitly quantify the rate at which accessory genes are integrated and lost. Consistent with theoretical expectations by Vos et al. (19), our results clearly suggest that the ratio of gene content changes to mutation was indeed very high at the narrow evolutionary scale analysed herein. Chromosome and plasmid gene substitutions per gene and per year were 7.00×10^−5^ and 1.75×10^−3^, respectively. Importantly, the genuine plasmid gene substitution rate may differ from our observation because the evolutionary history of horizontally transferred DNA may differ from that of the bacterial host. Here, we intend to determine the gene turnover rate of the bacterial host. Whether such a high gene turnover is being selected because it secures the access to rare but beneficial accessory genes remains debated (19, 55).

The observed high heterogeneity in gene turnover rates between subclades probably reflects the fact that numerous genes are gained simultaneously through plasmid integration. In the SWIO lineage, pCu^R^ is a good example of a recently acquired and successfully maintained mobile genetic element. However, the future trajectory of pXac39, a previously uncharacterized plasmid observed in a single strain, remains unknown and will depend on how it affects host fitness. Overall, plasmids appeared much more permeable to exogenous genetic material than chromosomes, with a gene turnover rate two orders of magnitude higher. Therefore, plasmids represent a privileged type of vehicle for accessory genes in *Xcc*, which is consistent with earlier network analyses (9).

### Concluding remarks

In this study, we reconstructed the genomic evolution of a lineage of the citrus pathogenic bacterium *Xcc* from its emergence in SWIO in the 19^th^ century to its global endemicity with epidemic waves nowadays. This lineage displayed a low SNP substitution rate, characteristic of monomorphic bacterium. In contrast to this apparently slow rate of evolution, we revealed the cardinal importance of genomic evolution through gene gain and loss. Plasmids played a key role in gene-based evolution: a drug-resistance plasmid spread among the SWIO lineage and pathogenicity-related plasmids exhibited extensive plasticity. By favouring intra-cellular recombination, *Tn3* transposons and IS*3* elements appear to promote genomic plasticity. Lastly, by describing chromosomal TALE genes for the first time in broad host range *Xcc* strains, our study also opens up a new line of investigation in the field of host/pathogen interaction.

## Materials and Methods

### DNA extraction and sequencing of bacterial strains

The dataset comprised whole genome sequences for 284 *Xcc* strains, including 210 strains from the South West Indian Ocean (SWIO; Figure 1, Supplementary Table S2). All the strains were sequenced using Illumina. Long-read Oxford Nanopore MinION sequencing was also performed on 13 selected strains. Copper-resistance phenotypes were determined previously (46). DNA sequencing and data processing are detailed in (56).

### SNP detection

We used a custom bioinformatics pipeline to obtain a filtered set of SNPs from the Illumina raw reads (Supplementary Figure S5). In short, after a quality control trimming step using Trimmomatic v. 0.36 (57), reads were aligned against the chromosome of *Xcc* strain IAPAR 306 (GenBank accession NC_003919.1) with BWA-MEM v. 0.7.15 (58). *Xcc* strains displayed a mean coverage of 232 (Table S2). Duplicated reads were removed using PicardTools MarkDuplicates v. 2.7. Indel realignment and SNP calling were performed using Freebayes 0.9.21-5 (59). SNPs were then filtered based on allele number, coverage, phred quality, allele frequency or genomic characteristics, such as SNP density or repeated genomic regions. In order to exclude SNPs found in regions that had undergone homologous recombination from the follow-up phylogenetic analyses, we searched for recombinant regions with ClonalFrameML (60) and RDP4 (61) using default parameters. Recombination analysis using ClonalframeML detected a 136kb recombinant region, comprising 233 SNPs in seven strains originating from Mali, Senegal, Bangladesh and India. The RDP4 analysis detected a 197kb (315 SNPs) recombinant region entirely overlapping the region detected with ClonalFrameML. These 315 SNPs were thus excluded from the global dataset for subsequent phylogenetic reconstruction and molecular clock analysis.

We first tested the adequacy of several models of molecular evolution with our SNP set. According to PartitionFinder v.2.1.1 and based on Bayesian Information Criterion, the model of evolution best fitting our dataset was a General Time-Reversible substitution model of evolution with variation among sites modelled with a discrete Gamma distribution and Invariant sites (GTR+G+I) (62). We reconstructed a Maximum Likelihood tree of the global dataset using RAxML v.8.2.9 (63). The presence of the temporal signal in the dataset was tested by computing the linear regression between sample age and root-to-tip distances at every internal node of the Maximum Likelihood tree (64). The SWIO clade root was the deepest node for which both the linear regression was statistically significant and the slope positive. The SWIO clade was, therefore, assumed to contain detectable amounts of evolutionary change, making it suitable for tip-dating inferences. A date-randomization test with 20 replicates was performed to confirm the presence of the temporal signal (65).

Tip-dating inference was then performed on the SWIO subset using BEAST v1.8.4 (30) with a GTR+G+I substitution model of evolution. We used an uncorrelated lognormal relaxed clock to account for rate variation among lineages. To minimize prior assumptions about demographic history, we first used an extended Bayesian skyline plot to integrate data over different coalescent histories. After inspecting the demographic reconstruction, an exponential growth was established as a best fit for the tree prior. Three independent chains were run for 100 000 000 steps and sampled every 10 000 steps. The first 1 000 samples (10%) were discarded as burn-in. Convergence to the stationary distribution and sufficient sampling and mixing were checked by inspecting posterior samples using Tracer v1.6 (effective sample size >200) (66). After combining the three runs, a maximum clade credibility tree was obtained with TreeAnnotator v1.10.2 (30).

### Core and accessory genome assignation

Our estimation of the gene content of each strain from the SWIO clade relied on a two-step approach. We first estimated the total SWIO homologous set of genes with a pipeline combining *de novo* assembly, gene prediction and gene clustering (Supplementary Figure S6). We then defined the gene content of individual strains. To validate our gene content analysis method, we first analysed three sequenced replicates of two strains (*Xcc* LH201 and *Xcc* LE50), both already sequenced using the long-read Pacific Biosciences RSII technology (Supplementary Figure S6). Besides assessing the error rate associated with our pipeline, the replicates were used to tune the mapping parameters that were to consider a gene present in a strain. Importantly, distinct bacterial cultures and DNA extractions were performed for each of the replicates. Therefore, our estimate will confound the genuine variation resulting from gene loss during culture and the variation associated with the bioinformatics pipeline. Lastly, we mapped each strain’s reads on the set of 5 046 homologous genes (Supplementary Table S2). We defined the gene content of individual strains using the parameters minimizing gene content variations between replicates (a gene was considered present in a strain if coverage was ≥20x over at least 60% of its length).

Genomic location of gene clusters to the chromosome or plasmids was informed by the available circularized Pacific Biosciences-sequenced reference genomes. Genomic location of unassigned gene clusters was defined based on the location of genes co-occurring on the same contigs. Functions were assigned to the gene clusters according to their amino-acid homology (using a 30% identity/30% length threshold) with known Clusters of Orthologous Groups (COGs) based on a BLASTx search.

In a second BEAST analysis using the previously inferred tree topology, a discrete model was used to reconstruct the ancestral states of gene presence/absence. Three independent chains were run as described above. Mean and 95% HPD gene substitution rate, gain rate and loss rate were obtained from the parameter distribution using a custom Perl script and the R HDInterval package (67). Rates of gene gain and loss were defined as the number of state changes (present or absent) per site per year, using the mean number of plasmid-borne and chromosomal genes per strain as the site number.

With the exception of the thirteen Minion-sequenced strains (Supplementary Table S1), the nature of our sequencing data prevented us from assembling closed, circularized plasmid genomes. However, using each strain’s gene content, we could assess each strain for the presence of genes previously identified on closed plasmids. High quality PacBio-based circular plasmid sequences were used as the genomic reference for each plasmid’s gene content and synteny: pLJ207-7.3 (GenBank accession CP018853.1, from SWIO strain *Xcc* LJ207-7, related to pXac47 (33)), pLH276.2 (GenBank accession CP018856.1, from SWIO strain *Xcc* LH276, related to pXac64 (28)), and the copper-resistance plasmid pCu^R^ (GenBank accession CP018859, from SWIO strain *Xcc* LH201 (24)).

## Data availability

The sequencing data generated in this study have been published in the NCBI GenBank repository under accession numbers listed in Supplementary Table S2, Additional File 1.

## Acknowledgements

We would like to express our thanks to the Plant Protection Platform (3P, IBISA) and to S. Javegny, K. Boyer and J. Hascoat for their helpful contribution. The European Regional Development Fund (ERDF contract GURDT I2016-1731-0006632), the Réunion Region, the French government, the French Agropolis Foundation (Labex Agro – Montpellier, E-SPACE project number 1504-004), ANSES and CIRAD provided financial support. We would like to thank INRAPE (the Union of the Comoros), FAREI (Mauritius) and NBA (Seychelles) for providing us with diseased citrus material. This work was supported by the CIRAD - UMR AGAP HPC Data Center of the South Green Bioinformatics platform (http://www.southgreen.fr/).

## Author contributions

D.R., O.P. and P.L. designed and conceived the study; C.B. processed the samples in the wet-lab; D.R. performed computational analyses with inputs from P.L. and A.R.; D.R., P.L. and O.P. wrote the manuscript with inputs from all co-authors.

