## Supplementary File 1 for "Time-calibrated genomic evolution of a monomorphic bacterium during its establishment as an endemic crop pathogen"

**Running title:** Genomic evolution of an emergent pathogenic bacterium

Damien Richard[a,b,c,#]

Olivier Pruvost[a]

François Balloux[d]

Claudine Boyer[a]

Adrien Rieux[a]

Pierre Lefeuvre[a]

[a] Cirad, UMR PVBMT, F-97410 St Pierre, Réunion, France.
[b] ANSES, Plant Health Laboratory, F-97410 St Pierre, Réunion, France.
[c] Université de la Réunion, UMR PVBMT, F-97490 St Denis, Réunion, France.
[d] UCL Genetics Institute, University College London, London WC1E 6BT, UK.

### Competing interests

The authors declare that they have no competing interests.

### Keywords

Genomic evolution; Dated phylogeny; Citrus canker; Gene turnover rate; SNP substitution rate

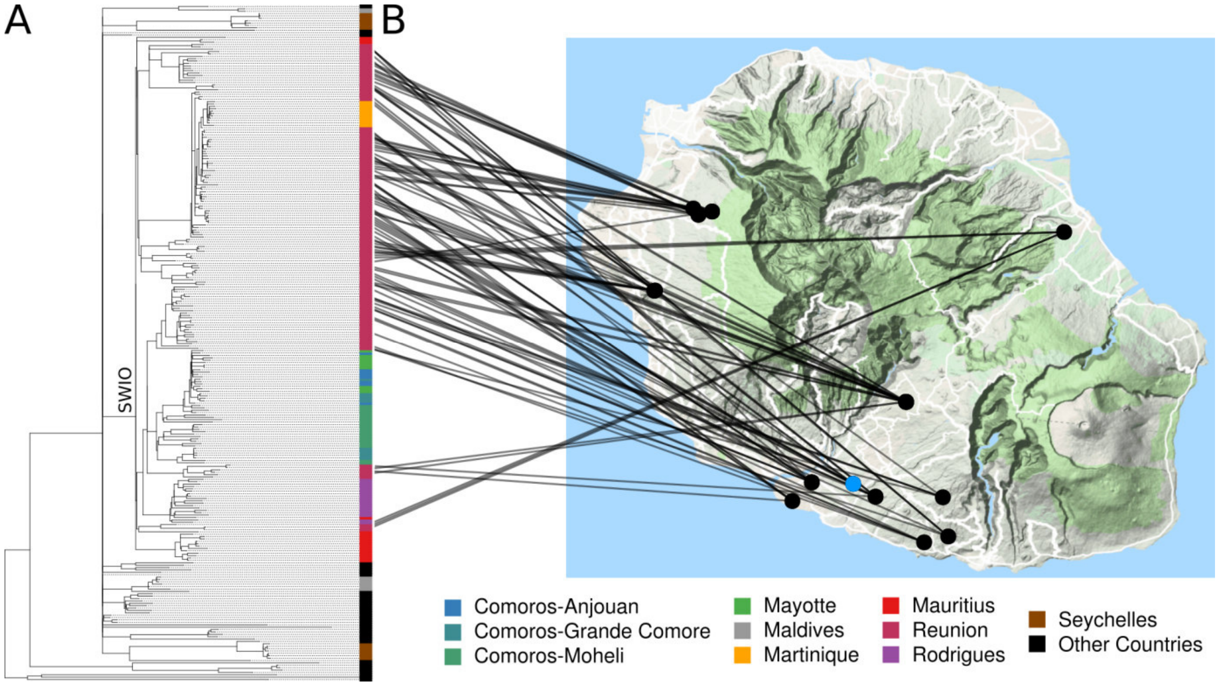


Figure S1. (A) Maximum likelihood tree of the global dataset. Boxes are coloured according to the strains' geographic origin (see legend). Node corresponding to SWIO clade root is annotated. (B) Map of Réunion linking the groves-replicates Réunion strains with the groves (black circles) and the nursery (the blue circle) where the strains were isolated.

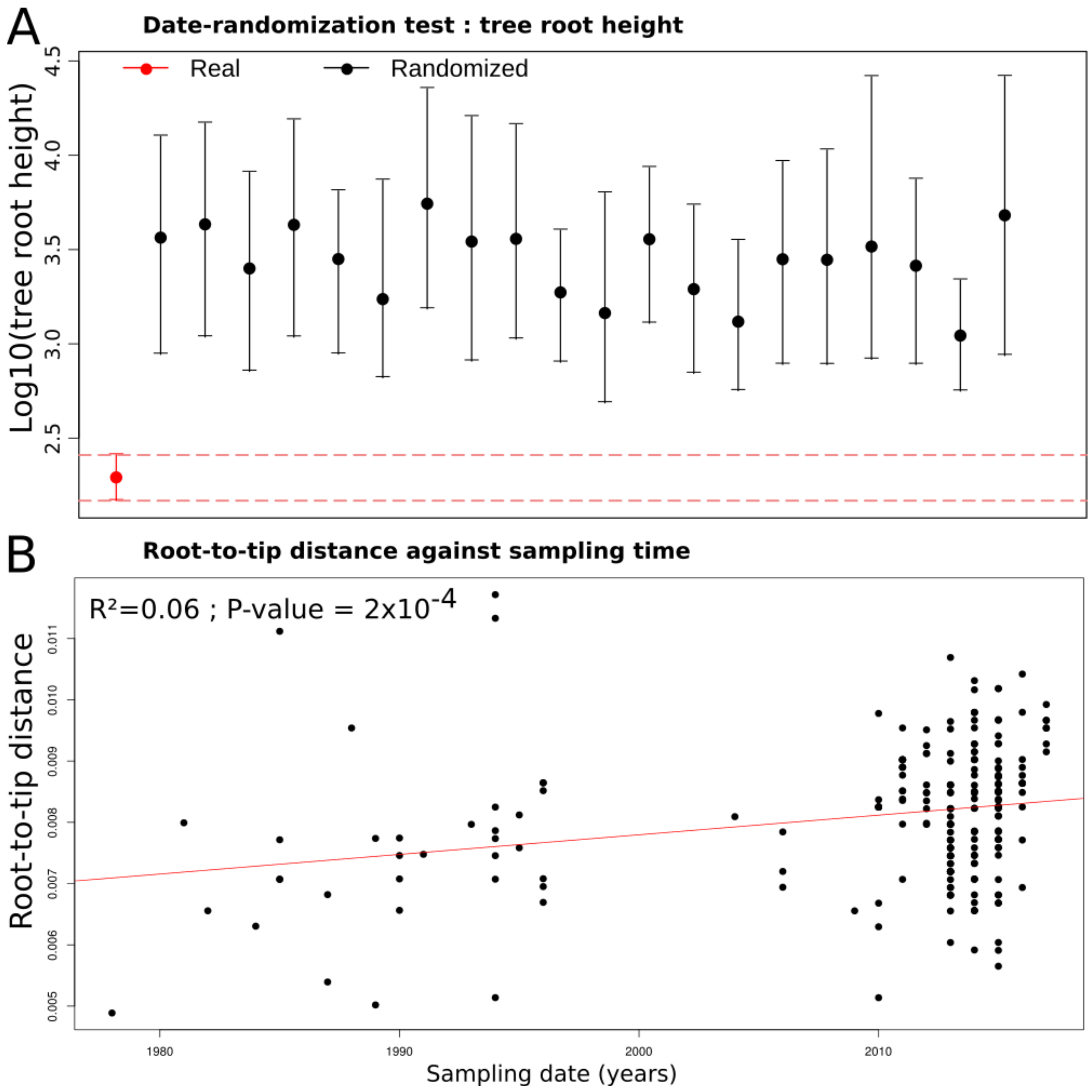


Figure S2. (A) Estimated SWIO tree root age and associated 95% HPD obtained from 20 BEAST analysis ran with randomized sample dates and from the real dataset. (B) Root-to-tip regression between root-to-tip distance and tip dates.

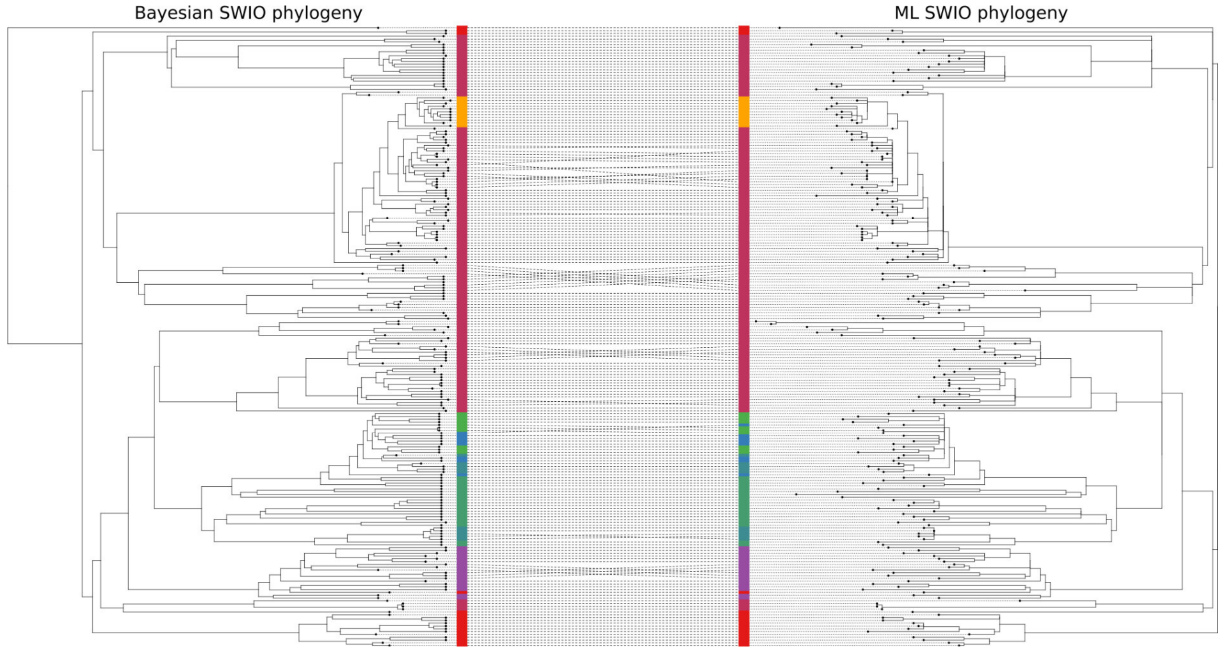


Figure S3. Comparison between the topologies of the Bayesian phylogeny (left) and the Maximum Likelihood phylogeny (right) of the SWIO strains. Tips are coloured according to sampling location, using the same colour code as in Figure 2.

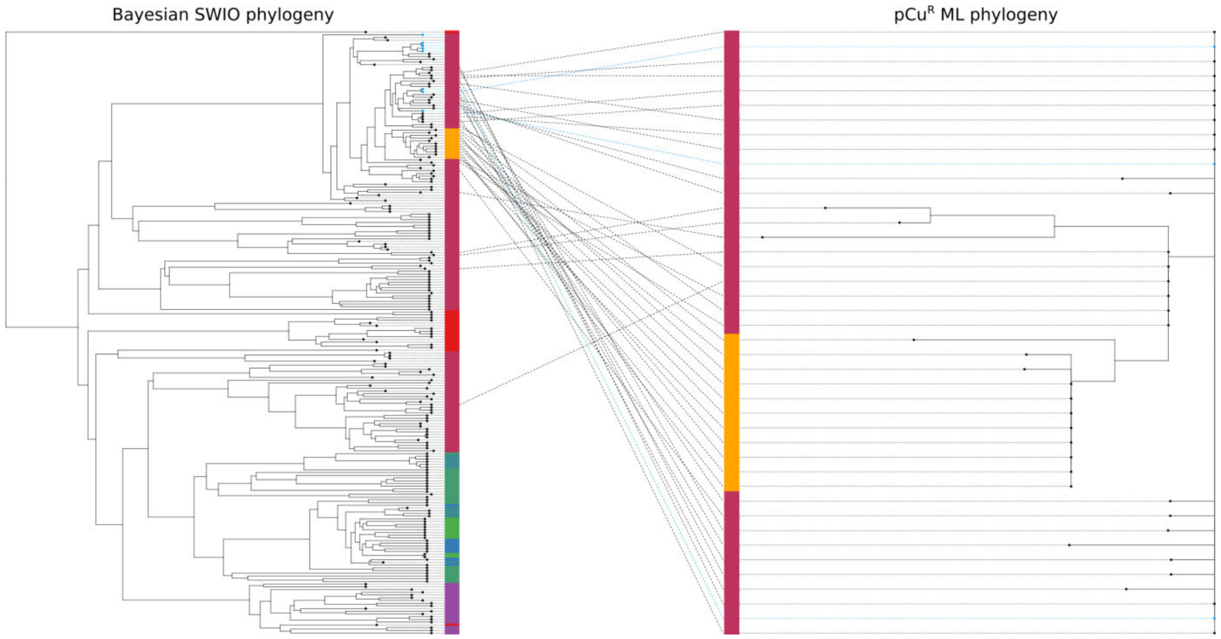


Figure S4. Bayesian phylogeny of the SWIO strains based on their chromosomal SNPs (left) and phylogeny of the copper-resistance plasmid present in some strains (right). Dotted lines join each strain with its plasmid. Blue circles at the tip of the branches of the phylogeny represent strains isolated in a Réunion nursery.

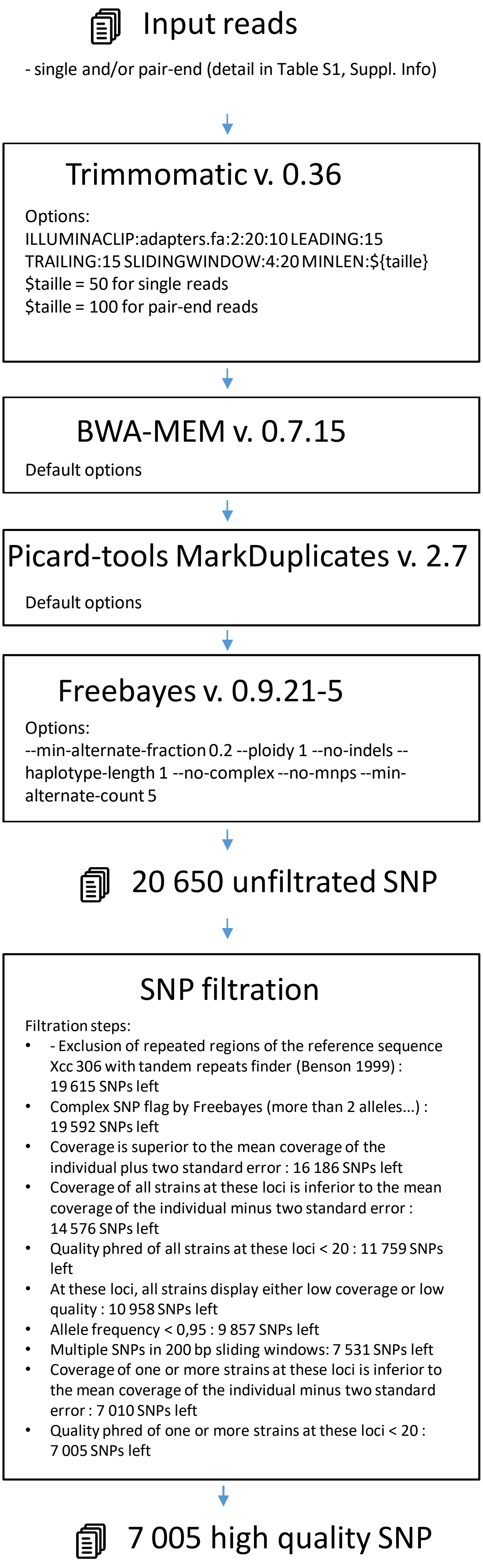


Figure S5. Schematics of the bioinformatics pipeline used for SNP inference.

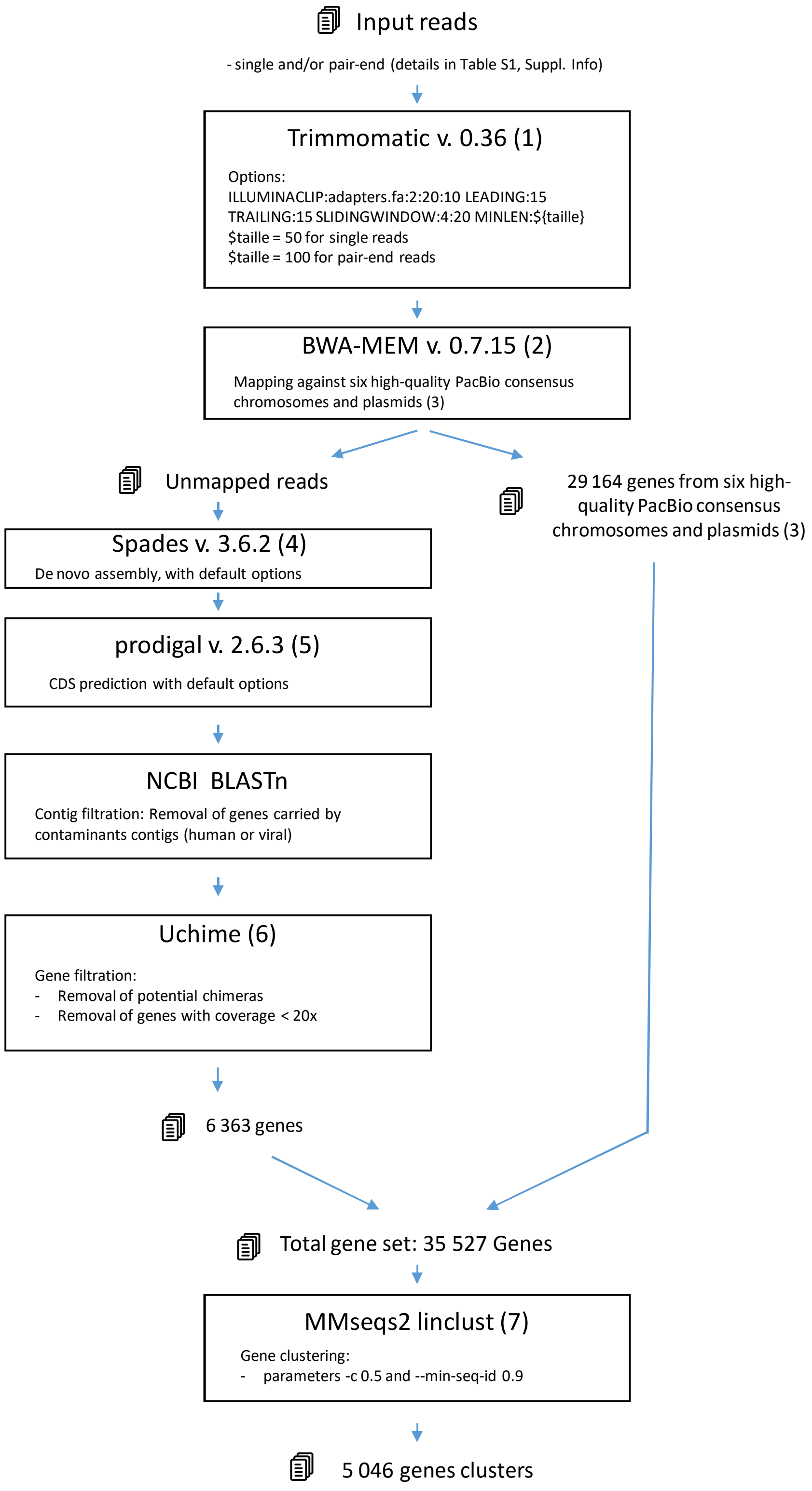


Figure S6. Schematics of the bioinformatics pipeline used to estimate the total SWIO homologous set of genes.

Table S1. Metrics of hybrid *de novo* assemblies (Nanopore MinION and Illumina) of 13 *Xcc* strains.

| Strain | Contig length | Suggested Circular | GenBank accession |
|---|---|---|---|
| JJ009-1 | 5153301 | yes | JAABBU000000000 |
| JJ009-1 | 89043 | yes | JAABBU000000000 |
| JJ009-1 | 39809 | yes | JAABBU000000000 |
| JJ009-1 | 19600 | yes | JAABBU000000000 |
| LK130-09 | 5152518 | yes | JAAAZW000000000 |
| LK130-09 | 66650 | yes | JAAAZW000000000 |
| LK130-09 | 22621 | no | JAAAZW000000000 |
| LK130-09 | 18354 | no | JAAAZW000000000 |
| LK136-05 | 5156448 | yes | JAABAG000000000 |
| LK136-05 | 48136 | yes | JAABAG000000000 |
| LK136-05 | 43807 | yes | JAABAG000000000 |
| LK136-05 | 22636 | yes | JAABAG000000000 |
| LK169-03 | 5151659 | yes | JAABFX000000000 |
| LK169-03 | 67511 | yes | JAABFX000000000 |
| LK169-03 | 44850 | yes | JAABFX000000000 |
| LK169-03 | 15612 | yes | JAABFX000000000 |
| LK145-14 | 5153497 | yes | JAAAZP000000000 |
| LK145-14 | 86788 | yes | JAAAZP000000000 |
| LK145-14 | 47146 | yes | JAAAZP000000000 |
| LK145-14 | 15762 | yes | JAAAZP000000000 |
| LK137-01 | 5096857 | yes | JAABAI000000000 |
| LK137-01 | 57839 | yes | JAABAI000000000 |
| LK137-01 | 56543 | yes | JAABAI000000000 |
| LK137-01 | 7969 | no | JAABAI000000000 |
| LL098-02 | 5158627 | yes | JAABGZ000000000 |
| LL098-02 | 74371 | no | JAABGZ000000000 |
| LL098-02 | 56043 | yes | JAABGZ000000000 |
| LL098-02 | 26975 | yes | JAABGZ000000000 |
| LL115-09 | 5157191 | yes | JAABGR000000000 |
| LL115-09 | 86709 | yes | JAABGR000000000 |
| LL115-09 | 46482 | no | JAABGR000000000 |
| LL115-09 | 15020 | no | JAABGR000000000 |
| LL174-05 | 5092686 | yes | JAABFG000000000 |
| LL174-05 | 71598 | no | JAABFG000000000 |
| LL174-05 | 59165 | no | JAABFG000000000 |
| LL174-05 | 46227 | no | JAABFG000000000 |
| LL174-05 | 26195 | no | JAABFG000000000 |
| LL174-05 | 23383 | no | JAABFG000000000 |
| LM089-20 | 5157648 | yes | JAABDL000000000 |
| LM089-20 | 89725 | yes | JAABDL000000000 |
| LM095-04 | 5109809 | yes | JAABIF000000000 |
| LM095-04 | 56722 | yes | JAABIF000000000 |
| LM095-04 | 46445 | yes | JAABIF000000000 |
| LM095-04 | 34348 | no | JAABIF000000000 |
| LM096-08 | 5107202 | yes | JAABIL000000000 |
| LM096-08 | 73016 | yes | JAABIL000000000 |
| LM096-08 | 21077 | no | JAABIL000000000 |

| | | | |
|---|---|---|---|
| LM096-08 | 5702 | no | JAABIL000000000 |
| LN006-18 | 5164630 | yes | JAABDQ000000000 |
| LN006-18 | 214873 | yes | JAABDQ000000000 |
| LN006-18 | 86747 | yes | JAABDQ000000000 |
| LN006-18 | 52388 | yes | JAABDQ000000000 |
| LN006-18 | 2466 | no | JAABDQ000000000 |

Table S2. Characteristics of all the sequenced bacterial strains of this study and metrics related to sequencing and steps of the bioinformatics pipeline applied on the dataset. Copper phenotype of the strains was abbreviated as S for copper-sensitivity, Rlab for copper-resistance due to the CopLAB system and Rabcd for copper-resistance due to the CopABCD system.

| | | | | | | | | Mapping on Xcc 306 for SNP calling | | De novo assembly using all reads | | De novo assembly using reads unmapped on pacbio genoms | | |
|---|---|---|---|---|---|---|---|---|---|---|---|---|---|---|
| Strain | Host | Country | Site number | Date | Copper phenotype | Nanopore MinION sequenced | SWIO | Mean coverage | Coverage std | De novo contig number | De novo contig length | De novo contig number | De novo contig length | De novo assembly GenBank accession |
| B25 | *C. hystrix* | Reunion | NA | 1987 | S | no | yes | 60 | 28 | 83 | 5166326 | 17 | 3936 | JAABGV000000000 |
| C20 | *C.* x *paradisi* | Reunion | NA | 1988 | S | no | yes | 75 | 29 | 82 | 5176962 | 15 | 3568 | JAAAYX000000000 |
| CFBP 1209 | *C. maxima* | Hong Kong | NA | 1963 | S | no | no | 99 | 38 | 87 | 5237899 | NA | NA | JAABCD000000000 |
| CFBP 2525 | *C.* x *limon* | New Zealand | NA | 1956 | S | no | no | 84 | 31 | 80 | 5171353 | NA | NA | JAABDC000000000 |
| CFBP 2853 | *Citrus* sp. | New Zealand | NA | 1958 | S | no | no | 75 | 28 | 78 | 5127192 | NA | NA | JAABDD000000000 |
| CFBP 2855 | *Citrus* sp. | Japan | NA | 1962 | S | no | no | 53 | 24 | 80 | 5249198 | NA | NA | JAABCK000000000 |
| CFBP 2857 | Citrus sp. | Fiji | NA | 1976 | S | no | no | 87 | 30 | 77 | 5173983 | NA | NA | JAABCC000000000 |
| CFBP 2865 | *C.* × *aurantiifc* | Brazil | NA | 1976 | S | no | no | 65 | 26 | 81 | 5131542 | NA | NA | JAAAZR000000000 |
| CFBP 2908 | *C. reticulata* | Brazil | NA | 1980 | S | no | no | 73 | 30 | 80 | 5197611 | NA | NA | JAAAZS000000000 |
| CFBP1814 | *C. maxima* | Reunion | NA | 1978 | S | no | yes | 46 | 51 | 83 | 5176025 | 6 | 15609 | JAABHX000000000 |
| CFBP2852 | *Citrus* sp. | India | NA | NA | S | no | no | 397 | 103 | 151 | 5268707 | NA | NA | CCWI00000000.1 |
| D02 | *C.* × *tangelo* | Reunion | NA | 1989 | S | no | yes | 38 | 48 | 101 | 5129123 | 11 | 2470 | JAABDI000000000 |
| D07 | *C.* × *unshiu* | Reunion | NA | 1989 | S | no | yes | 73 | 27 | 74 | 5161449 | 15 | 3555 | JAABGU000000000 |
| FDC1083 | *C. reticulata* | Brazil | NA | 1980 | S | no | no | 842 | 211 | 127 | 5215546 | NA | NA | CCVZ00000000.1 |
| FDC217 | *C.* x *sinensis* | Brazil | NA | 2003 | S | no | no | 866 | 203 | 133 | 5215799 | NA | NA | CCWY00000000.1 |
| JA159-1 | *C.* × *clementir* | Reunion | NA | 1981 | S | no | yes | 88 | 33 | 67 | 5174061 | 14 | 3168 | JAABGE000000000 |
| JB003-5 | *C.* × *aurantiifc* | Reunion | NA | 1982 | S | no | yes | 59 | 23 | 85 | 5193147 | 65 | 16390 | JAABEI000000000 |
| JJ009-04 | *C.* x *bergamia* | Mauritius | NA | 1985 | S | no | yes | 221 | 35 | 66 | 5142023 | 274 | 76477 | JAABHD000000000 |
| JJ009-1 | *P. trifoliata* x | Mauritius | NA | 1984 | S | yes[$] | yes | 90 | 32 | 79 | 5216258 | 20 | 44379 | JAABHE000000000 |
| JJ009-8 | *C.* x *sinensis* | Mauritius | NA | 1987 | S | no | yes | 80 | 28 | 78 | 5175822 | 14 | 3304 | JAABEK000000000 |
| JJ010-02 | *C.* × *aurantiifc* | Rodrigues | NA | 1985 | S | no | yes | 255 | 52 | 69 | 5217038 | 14 | 4887 | JAABAV000000000 |
| JJ010-4 | *C.* × *aurantiifc* | Rodrigues | NA | 1985 | S | no | yes | 70 | 31 | 86 | 5123551 | 19 | 18236 | JAABAW000000000 |
| JJ053-06 | *C. reticulata* | Taiwan | NA | 1977 | S | no | no | 88 | 35 | 89 | 5189352 | NA | NA | JAABJF000000000 |
| JJ10-1 | *C.* × *aurantiifc* | Rodrigues | NA | 1985 | S | no | yes | 171 | 54 | 258 | 5163969 | 0 | 0 | CDDV01000000 |
| JJ155 | Citrus sp. | Argentina | NA | 1977 | S | no | no | 56 | 22 | 80 | 5112812 | NA | NA | JAAAZL000000000 |
| JJ165 | *C. maxima* | India | NA | 1978 | S | no | no | 72 | 26 | 84 | 5199660 | NA | NA | JAABCF000000000 |
| JJ238-04 | *C.* × *aurantiifc* | Maldives | NA | 1987 | S | no | no | 637 | 136 | 69 | 5253532 | NA | NA | JAABCL000000000 |
| JJ238-06 | *C.* × *aurantiifc* | Maldives | NA | 1987 | S | no | no | 278 | 61 | 85 | 5266067 | NA | NA | JAABCM000000000 |
| JJ238-07 | *C.* × *aurantiifc* | Maldives | NA | 1987 | S | no | no | 237 | 49 | 69 | 5257582 | NA | NA | JAABCN000000000 |
| JJ238-08 | *C.* × *aurantiifc* | Maldives | NA | 1987 | S | no | no | 274 | 56 | 67 | 5247531 | NA | NA | JAABCO000000000 |
| JJ238-09 | *C.* × *aurantiifc* | Maldives | NA | 1987 | S | no | no | 219 | 52 | 68 | 5308367 | NA | NA | JAABCP000000000 |
| JJ238-10 | *C.* × *aurantiifc* | Maldives | NA | 1987 | S | no | no | 1031 | 236 | 142 | 5257926 | NA | NA | CCWC00000000.1 |
| JJ238-11 | *C.* × *aurantiifc* | Maldives | NA | 1987 | S | no | no | 644 | 124 | 83 | 5266873 | NA | NA | JAABCR000000000 |
| JJ238-16 | *C.* x *sinensis* | Pakistan | NA | 1988 | S | no | no | 79 | 31 | 79 | 5293249 | NA | NA | JAABDG000000000 |
| JK002-10 | *C.* × *aurantiifc* | Saudi Arabia | NA | 1988 | S | no | no | NA | NA | NA | NA | NA | NA | CCWV00000000.1 |
| JK004-04 | *C.* × *aurantiifc* | Maldives | NA | 1987 | S | no | no | 69 | 25 | 83 | 5243375 | NA | NA | JAABCS000000000 |
| JK101-1 | *C.* x *paradisi* | Argentina | NA | 1990 | S | no | no | 77 | 31 | 76 | 5198010 | NA | NA | JAAAZD000000000 |
| JK161 | *C.* x *sinensis* | Mauritius | NA | 1990 | S | no | yes | 235 | 55 | 68 | 5193220 | 141 | 55678 | JAABEL000000000 |
| JK167-1 | *C.* × *aurantiifc* | Mauritius | NA | 1990 | S | no | yes | 69 | 27 | 79 | 5191833 | 12 | 2765 | JAABFT000000000 |
| JK169 | *Citrus* sp. | Mauritius | NA | 1990 | S | no | yes | 199 | 40 | 57 | 5180573 | 334 | 92812 | JAABFJ000000000 |
| JK170 | *C.* × *aurantiifc* | Mauritius | NA | 1990 | S | no | yes | 196 | 46 | 75 | 5227752 | 150 | 54593 | JAABFU000000000 |
| JK4-1 | *Citrus* sp. | China | NA | unknov | S | no | no | 110 | 37 | 320 | 5184604 | NA | NA | CDMR01000000 |
| JM027-2 | *C.* x *limon* | Reunion | NA | 1991 | S | no | yes | 60 | 45 | 91 | 5168298 | 27 | 6415 | JAABHZ000000000 |
| JN564 | *Citrus* sp. | Comoros (Gra | NA | 1993 | S | no | yes | 77 | 24 | 87 | 5181155 | 0 | 0 | JAABAE000000000 |
| JP541 | *C.* x *paradisi* | Reunion | 10 | 1994 | S | no | yes | 337 | 77 | 68 | 5170667 | 2737 | 710798 | JAAAYY000000000 |
| JP608 | *C.* x *paradisi* | Reunion | 10 | 1994 | S | no | yes | 192 | 46 | 68 | 5186957 | 7 | 1578 | JAAAYZ000000000 |
| JP637-01 | *C. reticulata* | Reunion | NA | 1994 | S | no | yes | 69 | 28 | 82 | 5176134 | 13 | 2926 | JAABHY000000000 |
| JP668 | *C. hystrix* | Reunion | 12 | 1994 | S | no | yes | 211 | 49 | 61 | 5170427 | 3 | 582 | JAABFD000000000 |
| JP669 | *C. hystrix* | Reunion | 12 | 1994 | S | no | yes | 210 | 63 | 61 | 5169419 | 15 | 3993 | JAABFE000000000 |
| JP710-06 | *C.* x *paradisi* | Reunion | 10 | 1994 | S | no | yes | 227 | 54 | 73 | 5181066 | 7 | 1631 | JAAAZA000000000 |
| JP727 | *C.* x *paradisi* | Reunion | 10 | 1994 | S | no | yes | 440 | 114 | 61 | 5177641 | 258 | 70117 | JAAAZB000000000 |
| JP764-01 | *C.* x *paradisi* | Reunion | 10 | 1994 | S | no | yes | 195 | 57 | 75 | 5198544 | 6 | 1785 | JAABAO000000000 |
| JQ612-17 | *C. hystrix* | Reunion | 12 | 1995 | S | no | yes | 385 | 88 | 69 | 5157581 | 282 | 77675 | JAABFF000000000 |
| JQ613-01 | *C. hystrix* | Reunion | 12 | 1995 | S | no | yes | 101 | 33 | 71 | 5171988 | 25 | 6029 | JAABFG000000000 |
| JS502-01 | *C.* x *paradisi* | Reunion | 15 | 1996 | S | no | yes | 221 | 63 | 70 | 5208276 | 37 | 9644 | JAABAP000000000 |
| JS503-05 | *C.* x *paradisi* | Reunion | 15 | 1996 | S | no | yes | 233 | 58 | 69 | 5214097 | 5 | 1306 | JAABAQ000000000 |
| JS536-01 | *C.* x *paradisi* | Reunion | 15 | 1996 | S | no | yes | 259 | 67 | 71 | 5211276 | 12 | 3117 | JAABAR000000000 |
| JS538-02 | *C.* x *paradisi* | Reunion | 15 | 1996 | S | no | yes | 78 | 34 | 90 | 5212835 | 25 | 6005 | JAABAS000000000 |
| JS858-01 | *C.* x *paradisi* | Reunion | 15 | 1996 | S | no | yes | 92 | 35 | 83 | 5223091 | 13 | 3127 | JAABAT000000000 |
| JW160 | *C.* × *aurantiifc* | Bangladesh | NA | 2000 | S | no | no | 753 | 212 | 173 | 5249955 | NA | NA | CCWH00000000.1 |
| JZ092 | *C.* x *limon* | Seychelles | NA | 2003 | S | no | no | 90 | 35 | 94 | 5252340 | NA | NA | JAABIZ000000000 |
| JZ094 | *C.* x *limon* | Seychelles | NA | 2003 | S | no | no | 231 | 51 | 72 | 5171719 | NA | NA | JAABJA000000000 |
| LA087-2 | *Citrus* sp. | Comoros (Gra | NA | 2004 | S | no | yes | 62 | 21 | 79 | 5179575 | 18 | 4425 | JAABAF000000000 |
| LB100-1 | *P. trifoliata* x | Seychelles | NA | 2005 | S | no | no | 135 | 43 | 299 | 5243463 | NA | NA | CDAV01000000 |
| LB100-3 | *P. trifoliata* x | Seychelles | NA | 2005 | S | no | no | 98 | 41 | 84 | 5260633 | NA | NA | JAABIY000000000 |
| LC004-1 | *C. maxima* | Viet Nam | NA | 2006 | S | no | no | 90 | 54 | 82 | 5192383 | NA | NA | JAABIO000000000 |
| LC045-02 | *C.* × *aurantiifc* | Rodrigues | NA | 2006 | S | no | yes | 304 | 75 | 67 | 5219249 | 31 | 9119 | JAABAY000000000 |
| LC046 | *C.* × *aurantiifc* | Rodrigues | NA | 2006 | S | no | yes | 367 | 83 | 65 | 5210371 | 21 | 5578 | JAABAZ000000000 |
| LC048 | *C.* × *aurantiifc* | Rodrigues | NA | 2006 | S | no | yes | 90 | 32 | 86 | 5212523 | 24 | 18928 | JAABBA000000000 |
| LC80 | *C. reticulata* x | Mali | NA | 2006 | S | no | no | 855 | 296 | 157 | 5226129 | NA | NA | CCWJ00000000.1 |
| LD7-1 | *C.* × *aurantiifc* | Mali | NA | 2007 | S | no | no | 67 | 27 | 1559 | 5304045 | NA | NA | CDAL00000000.1 |
| LE116-1 | *C.* × *aurantiifc* | Mali | NA | 2008 | S | no | no | 96 | 37 | 1312 | 5324845 | NA | NA | CDHD01000000 |
| LG102 | *Citrus* sp. | Bangladesh | NA | 2006 | S | no | no | 113 | 57 | 1150 | 5286995 | NA | NA | CDAN01000000 |
| LG117 | *C.* × *aurantiifc* | Bangladesh | NA | 2009 | S | no | no | 122 | 38 | 338 | 5225477 | NA | NA | CDAX01000000 |
| LG136-04 | *C. hystrix* | Reunion | 11 | 2009 | S | no | yes | 73 | 29 | 82 | 5171760 | 37 | 9416 | JAABDO000000000 |
| LG97 | *C.* x *limon* | Bangladesh | NA | 2006 | S | no | no | 120 | 47 | 1124 | 5221084 | NA | NA | CDAK01000000 |
| LG98 | *C.* × *aurantiifc* | Bangladesh | NA | 2006 | S | no | no | 105 | 33 | 323 | 5151129 | NA | NA | CDBA01000000 |
| LH201 | *C. hystrix* | Reunion | 5 | 2010 | Rlab | no | yes | 111 | 38 | 67 | 5373883 | 16 | 3984 | JAABHV000000000 |
| LH212 | *C. hystrix* | Reunion | 5 | 2010 | S | no | yes | 248 | 65 | 72 | 5182077 | 9 | 2194 | JAABEM000000000 |
| LH221 | *C. hystrix* | Reunion | 5 | 2010 | S | no | yes | 198 | 46 | 71 | 5212091 | 3 | 423 | JAABEN000000000 |

| | | | | | | | | | | | | | | |
|---|---|---|---|---|---|---|---|---|---|---|---|---|---|---|
| LH238 | *C. hystrix* | Reunion | 8 | 2010 | S | no | yes | 290 | 66 | 70 | 5183711 | 329 | 92988 | JAABEO000000000 |
| LH240 | *C. hystrix* | Reunion | 8 | 2010 | S | no | yes | 205 | 47 | 59 | 5171325 | 2 | 534 | JAABEP000000000 |
| LH241 | *C. hystrix* | Reunion | 8 | 2010 | S | no | yes | 68 | 24 | 76 | 5164190 | 11 | 2573 | JAABEQ000000000 |
| LH276 | *C. reticulata* x | Reunion | NA | 2010 | Rlab | no | yes | 60 | 22 | 93 | 5414043 | 8 | 1605 | JAABGF000000000 |
| LH320 | *C. hystrix* | Reunion | 11 | 2010 | S | no | yes | 207 | 46 | 70 | 5182209 | 7 | 1779 | JAABER000000000 |
| LH322 | *C. hystrix* | Reunion | 11 | 2010 | S | no | yes | 305 | 68 | 68 | 5185742 | 374 | 114660 | JAABES000000000 |
| LH37-1 | *C. x paradisi* | Senegal | NA | 2010 | S | no | no | 120 | 38 | 417 | 5376274 | NA | NA | CDAS00000000.1 |
| LI070-01 | *C. hystrix* | Reunion | 6 | 2011 | Rlab | no | yes | 401 | 87 | 65 | 5397252 | 963 | 306718 | JAABDJ000000000 |
| LI070-02 | *C. hystrix* | Reunion | 6 | 2011 | Rlab | no | yes | 379 | 101 | 58 | 5384778 | 3736 | 972520 | JAABDK000000000 |
| LI070-03 | *C. hystrix* | Reunion | 6 | 2011 | Rlab | no | yes | 209 | 50 | 59 | 5383800 | 472 | 124614 | JAABDL000000000 |
| LI070-05 | *C. hystrix* | Reunion | 6 | 2011 | Rlab | no | yes | 322 | 74 | 62 | 5384245 | 2091 | 753393 | JAABDM000000000 |
| LI139-03 | *C. × aurantiifc* | Rodrigues | NA | 2011 | S | no | yes | 293 | 62 | 68 | 5227302 | 22 | 20541 | JAABBB000000000 |
| LI139-05 | *C. × aurantiifc* | Rodrigues | NA | 2011 | S | no | yes | 336 | 69 | 67 | 5214462 | 15 | 4295 | JAABBC000000000 |
| LI139-6 | *C. × aurantiifc* | Rodrigues | NA | 2011 | S | no | yes | 61 | 22 | 87 | 5217734 | 25 | 20011 | JAABBD000000000 |
| LI214-01 | *C. × aurantiifc* | Reunion | 1 (nurse | 2011 | S | no | yes | 309 | 68 | 62 | 5171239 | 56 | 13930 | JAABFW000000000 |
| LI214-02 | *C. × aurantiifc* | Reunion | 1 (nurse | 2011 | Rlab | no | yes | 221 | 53 | 58 | 5385187 | 945 | 285159 | JAABFX000000000 |
| LI214-05 | *C. × aurantiifc* | Reunion | 1 (nurse | 2011 | S | no | yes | 250 | 59 | 61 | 5173608 | 1322 | 382462 | JAABFY000000000 |
| LI214-07 | *C. × aurantiifc* | Reunion | 1 (nurse | 2011 | S | no | yes | 234 | 58 | 61 | 5171033 | 759 | 209427 | JAABFZ000000000 |
| LI214-09 | *C. × aurantiifc* | Reunion | 1 (nurse | 2011 | Rlab | no | yes | 266 | 61 | 62 | 5390743 | 413 | 118799 | JAABGA000000000 |
| LI214-10 | *C. × aurantiifc* | Reunion | 1 (nurse | 2011 | S | no | yes | 280 | 63 | 62 | 5171350 | 612 | 168865 | JAABGB000000000 |
| LI214-14 | *C. × aurantiifc* | Reunion | 1 (nurse | 2011 | S | no | yes | 384 | 83 | 64 | 5174662 | 2368 | 822916 | JAABGC000000000 |
| LI214-16 | *C. × aurantiifc* | Reunion | 1 (nurse | 2011 | Rlab | no | yes | 370 | 77 | 60 | 5391209 | 2368 | 861767 | JAABGD000000000 |
| LJ001-1 | *Citrus* sp. | Seychelles | NA | 2012 | S | no | no | 209 | 50 | 74 | 5254165 | NA | NA | JAABJE000000000 |
| LJ003-1 | *C. x sinensis* | Seychelles | NA | 2012 | S | no | no | 58 | 27 | 95 | 5243802 | NA | NA | JAABJB000000000 |
| LJ125-01 | *C. x sinensis* | Mayotte | NA | 2012 | S | no | yes | 277 | 60 | 66 | 5208879 | 12 | 18878 | JAABGJ000000000 |
| LJ207-7 | *C. hystrix* | Reunion | 5 | 2012 | Rlab | no | yes | 62 | 24 | 84 | 5408423 | 12 | 2753 | JAABDN000000000 |
| LJ225-01 | *C. x sinensis* | Mayotte | NA | 2012 | S | no | yes | 303 | 65 | 60 | 5166363 | 15 | 19602 | JAABET000000000 |
| LJ225-2 | *C. x sinensis* | Mayotte | NA | 2012 | S | no | yes | 317 | 69 | 61 | 5167280 | 21 | 21322 | JAABEU000000000 |
| LJ226-02 | *C. x sinensis* | Mayotte | NA | 2012 | S | no | yes | 324 | 71 | 65 | 5192769 | 9 | 2876 | JAABEV000000000 |
| LJ228-1 | *C. hystrix* | Mayotte | NA | 2012 | S | no | yes | 264 | 55 | 66 | 5192861 | 14 | 4142 | JAABHU000000000 |
| LJ229-06 | *C. reticulata* | Mayotte | NA | 2012 | S | no | yes | 80 | 31 | 75 | 5180062 | 18 | 4252 | JAABGI000000000 |
| LJ230-03 | *C. x sinensis* | Mayotte | NA | 2012 | S | no | yes | 457 | 99 | 64 | 5192915 | 868 | 270243 | JAABEW000000000 |
| LJ231-09 | *Citrus* sp. | Mayotte | NA | 2012 | S | no | yes | 317 | 67 | 66 | 5208693 | 21 | 21281 | JAABDB000000000 |
| LJ232-01 | *C. x sinensis* | Mayotte | NA | 2012 | S | no | yes | 255 | 55 | 66 | 5193645 | 11 | 2768 | JAABCZ000000000 |
| LJ232-05 | *C. x sinensis* | Mayotte | NA | 2012 | S | no | yes | 307 | 62 | 64 | 5195538 | 18 | 6996 | JAABDA000000000 |
| LK126-01 | *C. × aurantiifc* | Comoros (Anj | NA | 2013 | S | no | yes | 217 | 51 | 63 | 5194707 | 276 | 75286 | JAAAZW000000000 |
| LK126-02 | *C. × aurantiifc* | Comoros (Anj | NA | 2013 | S | no | yes | 229 | 52 | 66 | 5207629 | 1430 | 454048 | JAAAZX000000000 |
| LK126-03 | *C. × aurantiifc* | Comoros (Anj | NA | 2013 | S | no | yes | 259 | 58 | 71 | 5192174 | 150 | 40535 | JAAAZY000000000 |
| LK127-01 | *C. x sinensis* | Comoros (Anj | NA | 2013 | S | no | yes | 208 | 46 | 63 | 5189573 | 380 | 109623 | JAABAA000000000 |
| LK127-03 | *C. x sinensis* | Comoros (Anj | NA | 2013 | S | no | yes | 246 | 54 | 63 | 5183106 | 58 | 15551 | JAABAB000000000 |
| LK128-01 | *C. x sinensis* | Comoros (Anj | NA | 2013 | S | no | yes | 239 | 53 | 65 | 5211178 | 587 | 170401 | JAABAC000000000 |
| LK128-02 | *C. x sinensis* | Comoros (Anj | NA | 2013 | S | no | yes | 232 | 60 | 68 | 5210770 | 1066 | 336817 | JAABAD000000000 |
| LK129-03 | *C. × aurantiifc* | Comoros (Anj | NA | 2013 | S | no | yes | 271 | 59 | 68 | 5191972 | 949 | 283118 | JAAAZZ000000000 |
| LK130-04 | *C. × aurantiifc* | Comoros (Mol | NA | 2013 | S | no | yes | 246 | 51 | 65 | 5214094 | 16 | 19855 | JAABBI000000000 |
| LK130-06 | *C. × aurantiifc* | Comoros (Mol | NA | 2013 | S | no | yes | 272 | 58 | 71 | 5219585 | 39 | 25589 | JAABBJ000000000 |
| LK130-09 | *C. × aurantiifc* | Comoros (Mol | NA | 2013 | S | yes[$] | yes | 287 | 64 | 61 | 5174900 | 11 | 17737 | JAABBK000000000 |
| LK130-11 | *C. × aurantiifc* | Comoros (Mol | NA | 2013 | S | no | yes | 596 | 127 | 67 | 5211243 | 892 | 299825 | JAABBL000000000 |
| LK131-01 | *C. x sinensis* | Comoros (Mol | NA | 2013 | S | no | yes | 279 | 60 | 65 | 5192054 | 12 | 3864 | JAABBV000000000 |
| LK131-04 | *C. × aurantiifc* | Comoros (Mol | NA | 2013 | S | no | yes | 340 | 79 | 61 | 5192267 | 12 | 3649 | JAABBM000000000 |
| LK131-10 | *C. × aurantiifc* | Comoros (Mol | NA | 2013 | S | no | yes | 288 | 71 | 66 | 5197518 | 7 | 2323 | JAABBN000000000 |
| LK132-03 | *C. x sinensis* | Comoros (Mol | NA | 2013 | S | no | yes | 269 | 57 | 70 | 5211934 | 39 | 25638 | JAABBW000000000 |
| LK132-08 | *C. × aurantiifc* | Comoros (Mol | NA | 2013 | S | no | yes | 304 | 65 | 59 | 5158236 | 14 | 18462 | JAABBO000000000 |
| LK135-03 | *Citrus* sp. | Comoros (Mol | NA | 2015 | S | no | yes | 329 | 70 | 67 | 5200417 | 953 | 299174 | JAABCB000000000 |
| LK136-01 | *C. × aurantiifc* | Comoros (Mol | NA | 2013 | S | no | yes | 290 | 73 | 61 | 5174120 | 11 | 18027 | JAABBP000000000 |
| LK136-04 | *C. × aurantiifc* | Comoros (Mol | NA | 2013 | S | no | yes | 291 | 63 | 62 | 5174619 | 19 | 20875 | JAABBQ000000000 |
| LK136-05 | *C. × aurantiifc* | Comoros (Mol | NA | 2013 | S | yes[$] | yes | 272 | 63 | 63 | 5174174 | 24 | 21396 | JAABBR000000000 |
| LK136-08 | *C. × aurantiifc* | Comoros (Mol | NA | 2013 | S | no | yes | 283 | 62 | 58 | 5174920 | 9 | 17908 | JAABBS000000000 |
| LK137-01 | *C. x sinensis* | Comoros (Mol | NA | 2013 | S | yes[$] | yes | 332 | 68 | 64 | 5212479 | 12 | 18382 | JAABBX000000000 |
| LK137-02 | *C. × aurantiifc* | Comoros (Mol | NA | 2013 | S | no | yes | 242 | 53 | 68 | 5215470 | 23 | 21502 | JAABBT000000000 |
| LK141-03 | *C. x sinensis* | Comoros (Mol | NA | 2013 | S | no | yes | 247 | 53 | 70 | 5208698 | 13 | 18449 | JAABBY000000000 |
| LK141-09 | *C. x sinensis* | Comoros (Mol | NA | 2013 | S | no | yes | 477 | 107 | 62 | 5193668 | 1062 | 304776 | JAABBZ000000000 |
| LK141-15 | *C. x sinensis* | Comoros (Mol | NA | 2013 | S | no | yes | 235 | 52 | 68 | 5207961 | 15 | 19404 | JAABCA000000000 |
| LK142-04 | *C. × aurantiifc* | Comoros (Mol | NA | 2013 | S | no | yes | 343 | 85 | 64 | 5206274 | 1078 | 319173 | JAABBU000000000 |
| LK144-08 | *Citrus* sp. | Comoros (Gra | NA | 2013 | S | no | yes | 314 | 66 | 73 | 5215544 | 17 | 20702 | JAABAG000000000 |
| LK145-07 | *Citrus* sp. | Comoros (Gra | NA | 2013 | S | no | yes | 268 | 58 | 71 | 5216228 | 15 | 21353 | JAABAH000000000 |
| LK145-10 | *Citrus* sp. | Comoros (Gra | NA | 2013 | S | no | yes | 80 | 36 | 81 | 5207714 | 17 | 20494 | JAABAI000000000 |
| LK145-14 | *Citrus* sp. | Comoros (Gra | NA | 2013 | S | yes[$] | yes | 292 | 63 | 73 | 5223292 | 18 | 21854 | JAABAJ000000000 |
| LK148-03 | *Citrus* sp. | Comoros (Gra | NA | 2013 | S | no | yes | 239 | 53 | 74 | 5217790 | 12 | 20866 | JAABAK000000000 |
| LK169-01 | *C. hystrix* | Reunion | 11 | 2013 | S | no | yes | 71 | 25 | 80 | 5174041 | 15 | 3438 | JAABED000000000 |
| LK169-03 | *C. hystrix* | Reunion | 11 | 2013 | S | yes[$] | yes | 69 | 27 | 81 | 5112466 | 15 | 3663 | JAABEE000000000 |
| LK169-04 | *C. hystrix* | Reunion | 11 | 2013 | S | no | yes | 67 | 27 | 74 | 5113195 | 28 | 7201 | JAABEF000000000 |
| LK170-01 | *C. hystrix* | Reunion | 11 | 2013 | S | no | yes | 66 | 27 | 83 | 5117893 | 40 | 10053 | JAABEG000000000 |
| LK170-03 | *C. hystrix* | Reunion | 11 | 2013 | S | no | yes | 55 | 22 | 80 | 5124967 | 14 | 3434 | JAABEH000000000 |
| LK170-05 | *C. hystrix* | Reunion | 11 | 2013 | S | no | yes | 71 | 28 | 84 | 5176669 | 17 | 4073 | JAABFM000000000 |
| LK171-01 | *C. hystrix* | Reunion | 11 | 2013 | S | no | yes | 71 | 29 | 72 | 5019444 | 20 | 4786 | JAABFN000000000 |
| LK171-02 | *C. hystrix* | Reunion | 11 | 2013 | S | no | yes | 80 | 27 | 85 | 5170631 | 14 | 3217 | JAABFO000000000 |
| LK172-01 | *C. hystrix* | Reunion | 11 | 2013 | S | no | yes | 88 | 38 | 80 | 5172837 | 22 | 5375 | JAABFP000000000 |
| LK172-02 | *C. hystrix* | Reunion | 11 | 2013 | S | no | yes | 83 | 31 | 93 | 5173332 | 11 | 2715 | JAABFQ000000000 |
| LK173-01 | *C. hystrix* | Reunion | 11 | 2013 | S | no | yes | 85 | 30 | 83 | 5171806 | 10 | 2193 | JAABGK000000000 |
| LK173-02 | *C. hystrix* | Reunion | 11 | 2013 | S | no | yes | 83 | 29 | 91 | 5172729 | 14 | 3223 | JAABGL000000000 |
| LL068-04 | *Citrus* sp. | Comoros (Gra | NA | 2014 | S | no | yes | 282 | 60 | 63 | 5195115 | 19 | 5671 | JAABAL000000000 |
| LL068-06 | *Citrus* sp. | Comoros (Gra | NA | 2014 | S | no | yes | 300 | 64 | 63 | 5193172 | 17 | 4671 | JAABAM000000000 |
| LL068-07 | *Citrus* sp. | Comoros (Gra | NA | 2014 | S | no | yes | 299 | 60 | 59 | 5193522 | 54 | 14510 | JAABAN000000000 |
| LL074-04 | *C. x paradisi* | Martinique | NA | 2014 | Rlab | no | yes | 85 | 31 | 76 | 5381411 | 128 | 33406 | JAABHT000000000 |
| LL082-03 | *C. reticulata* x | Reunion | 3 | 2014 | S | no | yes | 194 | 43 | 69 | 5212255 | 4 | 923 | JAABGG000000000 |
| LL082-04 | *C. reticulata* x | Reunion | 3 | 2014 | Rlab | no | yes | 200 | 42 | 58 | 5385030 | 155 | 41356 | JAABGH000000000 |
| LL082-13 | *C. reticulata* x | Reunion | 3 | 2014 | S | no | yes | 195 | 48 | 71 | 5208760 | 4 | 930 | JAABDR000000000 |
| LL082-15 | *C. reticulata* x | Reunion | 3 | 2014 | Rlab | no | yes | 215 | 45 | 60 | 5384500 | 442 | 123768 | JAABDS000000000 |

| | | | | | | | | | | | | | | |
|---|---|---|---|---|---|---|---|---|---|---|---|---|---|---|
| LL082-16 | *C. reticulata* x | Reunion | 3 | 2014 | S | no | yes | 237 | 59 | 72 | 5211866 | 143 | 36961 | JAABDT000000000 |
| LL082-18 | *C. reticulata* x | Reunion | 3 | 2014 | Rlab | no | yes | 189 | 52 | 63 | 5382871 | 152 | 41208 | JAABDU000000000 |
| LL082-21 | *C. reticulata* x | Reunion | 3 | 2014 | S | no | yes | 520 | 129 | 68 | 5213547 | 1793 | 515737 | JAABDV000000000 |
| LL082-25 | *C. reticulata* x | Reunion | 3 | 2014 | S | no | yes | 229 | 54 | 69 | 5215977 | 4 | 895 | JAABDW000000000 |
| LL085-02 | *C.* x *limon* | Reunion | 4 | 2014 | Rlab | no | yes | 191 | 47 | 67 | 5376495 | 164 | 43410 | JAABIA000000000 |
| LL085-03 | *C.* x *limon* | Reunion | 4 | 2014 | S | no | yes | 252 | 56 | 67 | 5169149 | 12 | 2965 | JAABIB000000000 |
| LL087-01 | *C.* x *limon* | Reunion | 4 | 2014 | S | no | yes | 567 | 140 | 67 | 5212543 | 222 | 61965 | JAABIC000000000 |
| LL087-02 | *C.* x *limon* | Reunion | 4 | 2014 | S | no | yes | 213 | 51 | 65 | 5148305 | 5 | 1142 | JAABID000000000 |
| LL095-02 | *C. reticulata* x | Reunion | 14 | 2014 | S | no | yes | 204 | 47 | 71 | 5207156 | 6 | 1736 | JAABDX000000000 |
| LL095-08 | *C. reticulata* x | Reunion | 14 | 2014 | S | no | yes | 527 | 115 | 70 | 5207684 | 1214 | 434946 | JAABDY000000000 |
| LL095-20 | *C. reticulata* x | Reunion | 14 | 2014 | S | no | yes | 257 | 58 | 69 | 5212879 | 484 | 129922 | JAABEB000000000 |
| LL096-06 | *C.* x *limon* | Reunion | 16 | 2014 | S | no | yes | 216 | 50 | 70 | 5218100 | 2 | 392 | JAABIE000000000 |
| LL096-08 | *C.* x *limon* | Reunion | 16 | 2014 | S | no | yes | 592 | 128 | 64 | 5171070 | 489 | 136065 | JAABIF000000000 |
| LL096-09 | *C.* x *limon* | Reunion | 16 | 2014 | S | no | yes | 258 | 63 | 68 | 5206287 | 156 | 40067 | JAABIG000000000 |
| LL096-13 | *C.* x *limon* | Reunion | 16 | 2014 | S | no | yes | 204 | 59 | 69 | 5207985 | 5 | 1158 | JAABIH000000000 |
| LL098-02 | *C. reticulata* x | Reunion | 14 | 2014 | S | yes[$] | yes | 213 | 59 | 68 | 5208884 | 9 | 2122 | JAABEC000000000 |
| LL100-12 | *C. reticulata* x | Reunion | 14 | 2014 | S | no | yes | 186 | 57 | 70 | 5221538 | 5 | 1107 | JAABDZ000000000 |
| LL102-02 | *C.* x *limon* | Reunion | 16 | 2014 | S | no | yes | 237 | 56 | 71 | 5215830 | 5 | 1001 | JAABII000000000 |
| LL111-06 | *C.* x *sinensis* | Martinique | NA | 2014 | Rlab | no | yes | 65 | 23 | 82 | 5376769 | 16 | 3663 | JAABFS000000000 |
| LL115-09 | *C. reticulata* x | Reunion | 13 | 2014 | S | yes[$] | yes | 198 | 48 | 71 | 5214522 | 337 | 109003 | JAABEA000000000 |
| LL115-17 | *C. reticulata* x | Reunion | 13 | 2014 | S | no | yes | 208 | 50 | 72 | 5208391 | 3 | 470 | JAABGW000000000 |
| LL115-20 | *C. reticulata* x | Reunion | 13 | 2014 | S | no | yes | 226 | 54 | 72 | 5210033 | 2 | 400 | JAABGX000000000 |
| LL124-01 | *C.* x *sinensis* | Martinique | NA | 2014 | Rlab | no | yes | 115 | 37 | 72 | 5371489 | 34 | 8126 | JAABHL000000000 |
| LL131-03 | *C. reticulata* x | Reunion | 2 | 2014 | S | no | yes | 565 | 129 | 67 | 5180016 | 772 | 215902 | JAABGY000000000 |
| LL132-01 | *C. reticulata* x | Reunion | 2 | 2014 | S | no | yes | 262 | 58 | 61 | 5175940 | 597 | 187570 | JAABHG000000000 |
| LL162-01 | *C. reticulata* x | Reunion | 13 | 2014 | S | no | yes | 530 | 122 | 66 | 5146276 | 742 | 206627 | JAABHH000000000 |
| LL164-02 | *C. reticulata* x | Reunion | 13 | 2014 | S | no | yes | 273 | 68 | 72 | 5213273 | 5 | 1262 | JAABHI000000000 |
| LL174-01 | *C.* x *limon* | Reunion | 7 | 2014 | S | no | yes | 196 | 54 | 71 | 5152807 | 3 | 644 | JAABIJ000000000 |
| LL174-02 | *C.* x *limon* | Reunion | 7 | 2014 | S | no | yes | 331 | 75 | 70 | 5214176 | 613 | 182668 | JAABIK000000000 |
| LL174-04 | *C.* x *limon* | Reunion | 7 | 2014 | S | no | yes | 223 | 57 | 68 | 5185395 | 6 | 1288 | JAABIL000000000 |
| LL174-05 | *C.* x *limon* | Reunion | 7 | 2014 | S | yes[$] | yes | 211 | 62 | 71 | 5150773 | 6 | 1670 | JAABIM000000000 |
| LL175-01 | *C. reticulata* x | Reunion | 2 | 2014 | Rlab | no | yes | 265 | 65 | 62 | 5384285 | 1713 | 579885 | JAABHJ000000000 |
| LL186-5 | *C.* x *sinensis* | Reunion | NA | 2014 | S | no | yes | 88 | 31 | 83 | 5207388 | 15 | 3533 | JAABAU000000000 |
| LM053-06 | *C.* × *aurantiifc* | Mauritius | NA | 2015 | S | no | yes | 217 | 46 | 68 | 5185612 | 379 | 115522 | JAABFV000000000 |
| LM053-07 | *C.* × *aurantiifc* | Mauritius | NA | 2015 | S | no | yes | 592 | 127 | 64 | 5189495 | 979 | 284959 | JAABGP000000000 |
| LM054-06 | *C.* x *sinensis* | Mauritius | NA | 2015 | S | no | yes | 270 | 68 | 66 | 5186246 | 1062 | 331817 | JAABFH000000000 |
| LM054-17 | *C.* x *sinensis* | Mauritius | NA | 2015 | S | no | yes | 202 | 53 | 66 | 5186208 | 214 | 61867 | JAABFI000000000 |
| LM055-08 | *C. reticulata* : | Mauritius | NA | 2015 | S | no | yes | 52 | 22 | 81 | 5169048 | 10 | 2301 | JAABHC000000000 |
| LM057-04 | *C.* x *meyeri* | Mauritius | NA | 2015 | S | no | yes | 207 | 57 | 66 | 5187848 | 114 | 31613 | JAABHF000000000 |
| LM057-14 | *C.* x *meyeri* | Mauritius | NA | 2015 | S | no | yes | 198 | 43 | 65 | 5183873 | 589 | 167623 | JAABGM000000000 |
| LM057-15 | *C.* x *meyeri* | Mauritius | NA | 2015 | S | no | yes | 211 | 44 | 68 | 5178922 | 468 | 127995 | JAABEJ000000000 |
| LM069-01 | *C.* × *aurantiifc* | Mauritius | NA | 2015 | S | no | yes | 231 | 49 | 61 | 5172128 | 284 | 76459 | JAABGQ000000000 |
| LM070 | *C.* × *aurantiifc* | Mauritius | NA | 2015 | S | no | yes | 211 | 47 | 62 | 5166844 | 251 | 68102 | JAABHB000000000 |
| LM088-18 | *C. hystrix* | Reunion | 5 | 2015 | S | no | yes | 239 | 64 | 71 | 5182336 | 4 | 923 | JAABFK000000000 |
| LM088-20 | *C. hystrix* | Reunion | 5 | 2015 | Rlab | no | yes | 68 | 27 | 83 | 5387923 | 14 | 3432 | JAABFL000000000 |
| LM088-25 | *C. hystrix* | Reunion | 5 | 2015 | S | no | yes | 255 | 61 | 57 | 5173954 | 160 | 44026 | JAABDP000000000 |
| LM088-28 | *C. hystrix* | Reunion | 5 | 2015 | Rlab | no | yes | 253 | 60 | 66 | 5376335 | 843 | 237888 | JAABDQ000000000 |
| LM088-37 | *C. hystrix* | Reunion | 5 | 2015 | Rlab | no | yes | 210 | 48 | 59 | 5376383 | 344 | 96468 | JAABEX000000000 |
| LM088-40 | *C. hystrix* | Reunion | 5 | 2015 | S | no | yes | 213 | 58 | 62 | 5169790 | 3 | 543 | JAABEY000000000 |
| LM088-42 | *C. hystrix* | Reunion | 5 | 2015 | Rlab | no | yes | 243 | 57 | 65 | 5387984 | 637 | 169175 | JAABEZ000000000 |
| LM089-02 | *C. reticulata* x | Reunion | 2 | 2015 | Rlab | no | yes | 63 | 24 | 87 | 5402086 | 96 | 24413 | JAABGZ000000000 |
| LM089-06 | *C. reticulata* x | Reunion | 2 | 2015 | S | no | yes | 216 | 52 | 63 | 5166695 | 2 | 314 | JAABHA000000000 |
| LM089-20 | *C. reticulata* x | Reunion | 2 | 2015 | S | yes[$] | yes | 213 | 60 | 62 | 5166959 | 3 | 589 | JAABHK000000000 |
| LM089-41 | *C. reticulata* x | Reunion | 2 | 2015 | S | no | yes | 223 | 64 | 64 | 5166765 | 5 | 1228 | JAABGR000000000 |
| LM089-42 | *C. reticulata* x | Reunion | 2 | 2015 | Rlab | no | yes | 201 | 50 | 60 | 5384759 | 315 | 86559 | JAABGS000000000 |
| LM090-02 | *Citrus* sp. | Martinique | NA | 2015 | Rlab | no | yes | 105 | 35 | 68 | 5374377 | 74 | 18043 | JAABGO000000000 |
| LM095-04 | *C.* × *aurantiifc* | Rodrigues | NA | 2015 | S | yes[$] | yes | 268 | 64 | 71 | 5145603 | 18 | 4844 | JAABBE000000000 |
| LM095-05 | *C.* × *aurantiifc* | Rodrigues | NA | 2015 | S | no | yes | 250 | 59 | 61 | 5221945 | 36 | 24724 | JAABBF000000000 |
| LM095-07 | *C.* × *aurantiifc* | Rodrigues | NA | 2015 | S | no | yes | 296 | 65 | 64 | 5230681 | 12 | 18210 | JAABBG000000000 |
| LM095-11 | *C.* × *aurantiifc* | Rodrigues | NA | 2015 | S | no | yes | 212 | 48 | 68 | 5172835 | 7 | 17030 | JAABBH000000000 |
| LM095-14 | *C.* × *aurantiifc* | Rodrigues | NA | 2015 | S | no | yes | 307 | 76 | 62 | 5177716 | 10 | 2466 | JAABCU000000000 |
| LM096-03 | *C.* × *aurantiifc* | Rodrigues | NA | 2015 | S | no | yes | 256 | 59 | 69 | 5160121 | 18 | 4619 | JAABCV000000000 |
| LM096-08 | *C.* × *aurantiifc* | Rodrigues | NA | 2015 | S | yes[$] | yes | 388 | 86 | 65 | 5116784 | 31 | 10654 | JAABCW000000000 |
| LM096-09 | *C.* × *aurantiifc* | Rodrigues | NA | 2015 | S | no | yes | 288 | 68 | 69 | 5171067 | 101 | 26312 | JAABCX000000000 |
| LM097-01 | *C.* × *aurantiifc* | Rodrigues | NA | 2015 | S | no | yes | 282 | 64 | 65 | 5226260 | 14 | 18757 | JAABCY000000000 |
| LM121-01 | *C.* x *limon* | Reunion | 4 | 2015 | Rlab | no | yes | 87 | 33 | 76 | 5380803 | 15 | 3615 | JAABIN000000000 |
| LM121-02 | *C.* x *limon* | Reunion | 4 | 2015 | Rlab | no | yes | 312 | 76 | 62 | 5383880 | 539 | 148070 | JAAAYI000000000 |
| LM121-31 | *C.* x *limon* | Reunion | 4 | 2015 | S | no | yes | 234 | 60 | 69 | 5205127 | 8 | 2085 | JAAAYJ000000000 |
| LM121-41 | *C.* x *limon* | Reunion | 4 | 2015 | Rlab | no | yes | 327 | 72 | 63 | 5383648 | 612 | 170449 | JAAAYK000000000 |
| LM121-42 | *C.* x *limon* | Reunion | 4 | 2015 | Rlab | no | yes | 184 | 42 | 62 | 5383613 | 531 | 144918 | JAAAYL000000000 |
| LM158 | *C.* x *sinensis* | Argentina | NA | 2013 | Rabcd | no | no | 72 | 26 | 85 | 5401423 | NA | NA | JAAAZH000000000 |
| LM169 | *C.* x *paradisi* | Argentina | NA | 2005 | Rlab | no | no | 69 | 23 | 81 | 5418449 | NA | NA | JAAAZE000000000 |
| LM180 | *C.* x *paradisi* | Argentina | NA | 2003 | Rlab | yes[$] | no | 76 | 28 | 78 | 5384936 | NA | NA | JAAAZF000000000 |
| LM184 | *C.* x *limon* | Argentina | NA | 2013 | S | no | no | 74 | 27 | 75 | 5201330 | NA | NA | JAAAZC000000000 |
| LM198 | *C.* x *sinensis* | Argentina | NA | 2015 | S | no | no | 57 | 21 | 74 | 5198241 | NA | NA | JAAAZI000000000 |
| LM199 | *C.* x *sinensis* | Argentina | NA | 2015 | Rabcd | no | no | 83 | 29 | 81 | 5392132 | NA | NA | JAAAZJ000000000 |
| LM205 | *C.* x *paradisi* | Argentina | NA | 2010 | Rlab | no | no | 73 | 28 | 82 | 5413242 | NA | NA | JAAAZG000000000 |
| LM229 | *C.* x *sinensis* | Argentina | NA | 2012 | Rabcd | no | no | 90 | 31 | 84 | 5392621 | NA | NA | JAAAZK000000000 |
| LM358-02 | *C.* x *limon* | Reunion | 9 | 2015 | S | no | yes | 576 | 129 | 68 | 5184610 | 1973 | 584378 | JAAAYM000000000 |
| LM358-08 | *C.* x *limon* | Reunion | 9 | 2015 | S | no | yes | 333 | 81 | 68 | 5152527 | 51 | 13341 | JAAAYN000000000 |
| LM358-13 | *C.* x *limon* | Reunion | 9 | 2015 | S | no | yes | 231 | 57 | 71 | 5187169 | 6 | 1398 | JAAAYO000000000 |
| LM358-18 | *C.* x *limon* | Reunion | 9 | 2015 | S | no | yes | 206 | 53 | 65 | 5181284 | 4 | 906 | JAAAYP000000000 |
| LM358-22 | *C.* x *limon* | Reunion | 9 | 2015 | S | no | yes | 274 | 63 | 69 | 5181248 | 3 | 632 | JAAAYQ000000000 |
| LMG9322 | *C.* × *aurantiifc* | USA (Florida) | NA | 1986 | S | no | no | 746 | 201 | 1143 | 5107574 | NA | NA | JPYD00000000.1 |
| LN003-10 | *C. hystrix* | Reunion | 11 | 2016 | S | no | yes | 225 | 57 | 72 | 5184907 | 7 | 1927 | JAABFA000000000 |
| LN003-17 | *C. hystrix* | Reunion | 11 | 2016 | S | no | yes | 197 | 50 | 70 | 5183131 | 6 | 1418 | JAABFB000000000 |
| LN005-04 | *C. hystrix* | Reunion | 8 | 2016 | S | no | yes | 232 | 54 | 61 | 5169740 | 3 | 644 | JAABFC000000000 |
| LN005-07 | *C. hystrix* | Reunion | 8 | 2016 | S | no | yes | 207 | 50 | 63 | 5165628 | 415 | 130517 | JAABHW000000000 |

| | | | | | | | | | | | | | | |
|---|---|---|---|---|---|---|---|---|---|---|---|---|---|---|
| LN006-01 | *C. reticulata* x | Reunion | 2 | 2016 | Rlab | no | yes | 234 | 64 | 64 | 5389774 | 378 | 104564 | JAABGT000000000 |
| LN006-18 | *C. reticulata* x | Reunion | 2 | 2016 | Rlab | yes$ | yes | 178 | 56 | 72 | 5419891 | 238 | 64355 | JAABDH000000000 |
| LN007-02 | *C.* x *limon* | Reunion | 7 | 2016 | S | no | yes | 240 | 63 | 68 | 5172990 | 8 | 1819 | JAAAYR000000000 |
| LN007-03 | *C.* x *limon* | Reunion | 7 | 2016 | S | no | yes | 278 | 66 | 67 | 5188147 | 1549 | 470298 | JAAAYS000000000 |
| LN007-05 | *C.* x *limon* | Reunion | 7 | 2016 | Rlab | no | yes | 208 | 61 | 64 | 5390093 | 711 | 199934 | JAAAYT000000000 |
| LN007-07 | *C.* x *limon* | Reunion | 7 | 2016 | Rlab | no | yes | 222 | 49 | 64 | 5383947 | 127 | 33710 | JAAAYU000000000 |
| LN007-09 | *C.* x *limon* | Reunion | 7 | 2016 | Rlab | no | yes | 221 | 62 | 64 | 5389878 | 389 | 105919 | JAAAYV000000000 |
| LN007-10 | *C.* x *limon* | Reunion | 7 | 2016 | Rlab | no | yes | 245 | 63 | 64 | 5394877 | 83 | 19419 | JAAAYW000000000 |
| LP027-03 | *C.* × *aurantiifc* | Seychelles | NA | 2017 | S | no | no | 204 | 46 | 66 | 5166247 | NA | NA | JAABIR000000000 |
| LP027-05 | *C.* × *aurantiifc* | Seychelles | NA | 2017 | S | no | no | 253 | 56 | 70 | 5165276 | NA | NA | JAABIS000000000 |
| LP027-13 | *C.* × *aurantiifc* | Seychelles | NA | 2017 | S | no | no | 222 | 49 | 71 | 5163061 | NA | NA | JAABIT000000000 |
| LP028-03 | *C.* × *aurantiifc* | Seychelles | NA | 2017 | S | no | no | 211 | 48 | 69 | 5169346 | NA | NA | JAABIU000000000 |
| LP028-05 | *C.* × *aurantiifc* | Seychelles | NA | 2017 | S | no | no | 227 | 52 | 70 | 5168495 | NA | NA | JAABIV000000000 |
| LP028-06 | *C.* × *aurantiifc* | Seychelles | NA | 2017 | S | no | no | 224 | 50 | 71 | 5169894 | NA | NA | JAABIW000000000 |
| LP029-13 | *C.* x *sinensis* | Seychelles | NA | 2017 | S | no | no | 659 | 155 | 74 | 5256328 | NA | NA | JAABJC000000000 |
| LP029-15 | *C.* x *sinensis* | Seychelles | NA | 2017 | S | no | no | 215 | 52 | 69 | 5252500 | NA | NA | JAABJD000000000 |
| LP030-1 | *C.* x *latifolia* | Martinique | NA | 2017 | Rlab | no | yes | 248 | 49 | 62 | 5381949 | 124 | 35052 | JAABHP000000000 |
| LP031-1 | *C.* x *sinensis* | Martinique | NA | 2017 | Rlab | no | yes | 257 | 54 | 65 | 5385235 | 215 | 60355 | JAABGN000000000 |
| LP032-1 | *C.* × *aurantiifc* | Martinique | NA | 2017 | Rlab | no | yes | 274 | 57 | 64 | 5382678 | 441 | 123131 | JAABHO000000000 |
| LP033-1 | *C.* x *paradisi* | Martinique | NA | 2017 | Rlab | no | yes | 243 | 64 | 63 | 5381639 | 595 | 167373 | JAABFR000000000 |
| LP034-1 | *C.* x *latifolia* | Martinique | NA | 2017 | Rlab | no | yes | 238 | 54 | 63 | 5384153 | 581 | 172143 | JAABHQ000000000 |
| LP035-1 | *C.* x *latifolia* | Martinique | NA | 2017 | Rlab | no | yes | 200 | 43 | 59 | 5383539 | 546 | 152498 | JAABHR000000000 |
| LP036-1 | *C.* x *latifolia* | Martinique | NA | 2017 | Rlab | no | yes | 542 | 135 | 63 | 5394028 | 419 | 122358 | JAABHS000000000 |
| NCPPB 211 | *Citrus* sp. | India | NA | 1948 | S | no | no | 64 | 24 | 98 | 5187563 | NA | NA | JAABCI000000000 |
| NCPPB 226 | *Citrus* sp. | New Zealand | NA | 1949 | S | no | no | 74 | 28 | 80 | 5123687 | NA | NA | JAABDE000000000 |
| NCPPB3562 | *C.* x *limon* | India | NA | 1988 | S | no | no | 803 | 240 | 198 | 5512184 | NA | NA | CCXZ01000000 |
| NCPPB3610 | *P. trifoliata* | India | NA | 1988 | S | no | no | 93 | 35 | 1189 | 5151260 | NA | NA | CDAO00000000.1 |
| NCPPB3612 | *C.* × *aurantiifc* | India | NA | 1988 | S | no | no | 120 | 42 | 426 | 5319808 | NA | NA | CDAQ00000000.1 |
| NCPPB406 | *Citrus* sp. | New Zealand | NA | 1957 | S | no | no | 75 | 25 | 82 | 5172395 | NA | NA | JAABDF000000000 |

$ Strains also sequenced using MinIon Nanopore, see supplementary Table S1 for details.
